# Genozip Dual-Coordinate VCF format enables efficient genomic analyses and alleviates liftover limitations

**DOI:** 10.1101/2022.07.17.500374

**Authors:** Divon Lan, Gludhug Purnomo, Ray Tobler, Yassine Souilmi, Bastien Llamas

## Abstract

We introduce Dual Coordinate VCF (DVCF), a file format that records genomic variants against two different reference genomes simultaneously and is fully compliant with the current VCF specification. As implemented in the Genozip platform, DVCF enables bioinformatics pipelines to seamlessly operate across two coordinate systems by leveraging the system most advantageous to each pipeline step, simplifying bioinformatics workflows and reducing file generation and associated data storage burden. Moreover, our benchmarking of Genozip DVCF shows that it produces more complete, less erroneous, and less biased translations across coordinate systems than two widely used alternative tools (i.e., LiftoverVcf and CrossMap).

**Availability and Implementation:** An open source (GPL) version of Genozip containing DVCF functionality but not compression functionality, and which includes scripts for reproducing the benchmarks presented here, is available at https://github.com/divonlan/dvcf. Documentation is available at https://genozip.com/dvcf.

## Background

Genomic sequencing and assembly technologies continue to evolve at a rapid pace, enabling the creation of new and more accurate reference genomes for various species [1]. While improved human reference genomes are welcomed by researchers and clinicians, updated assemblies inevitably result in altered coordinates that can hinder their adoption as it is common that legacy datasets and bioinformatics software required for some analyses often still use the older coordinate system. Accordingly, for species such as humans where multiple reference genomes (or versions thereof) are available, analytical pipelines often need to alternate between two coordinate systems to accommodate these limitations.

To facilitate workflows involving data mapped against different reference genomes, software such as CrossMap [2] and GATK LiftoverVcf [3] translate genomic coordinates across the assemblies using a chain file [4]. Variants, typically encoded in a VCF (Variant Call Format) [5] file, are then lifted over from one coordinate system to the other, resulting in file duplication (i.e., one for each coordinate system). The liftover step is typically lossy, with variants discarded due their coordinates lacking alignment in the chain file as well as limitations of the liftover software used [1, 6]. Moreover, errors may be introduced due to incorrect chain file mapping of variants as well as incorrect annotation conversion, along with the introduction of potential biases due to the concentration of discarded variants in certain genomic regions.

### Description and Results

Here we introduce an implementation of the *Dual Coordinate VCF* (DVCF) [7] format in the Genozip platform [8, 9], an extensible compression software. DVCF is an extension to the standard VCF format compliant with the VCF 4.3 specification, that includes variants with coordinates pertaining to two different genome assemblies simultaneously (Fig. S1). Using DVCF files, researchers can alternate between coordinate systems according to their needs – without creating duplicate VCF files – thereby reducing workflow complexity and alleviating demands on time, computational resources, and disk storage burden (the latter being further improved by Genozip’s efficient data compression algorithms; Tables S16-17). Importantly, the DVCF file format is independent of its implementation in Genozip, allowing its implementation in any relevant bioinformatics software, thereby maintaining interoperability between tools.

We refer to the process of converting a VCF file to the DVCF format as *lifting*. In order to lift a VCF file to DVCF, Genozip requires three inputs: two reference files – one defining the *Primary* coordinates as used in the input VCF, and the other defining the *Luft* coordinates to be lifted over (“*luft*” is a neologism representing an alternative past-participle of “lift”) – and a chain file defining alignments between the two reference assemblies. Once a DVCF file is generated, users can render it in either *Primary* or *Luft* coordinates by using Genozip’s `genocat` command (see Figs. 1A, S2). *Cross-rendering* consists of losslessly re-arranging the information in the DVCF file to change the coordinate system in which the variants are represented. This allows users to seamlessly alternate between coordinate systems according to the particulars of their bioinformatic workflow.

**Figure 1.**
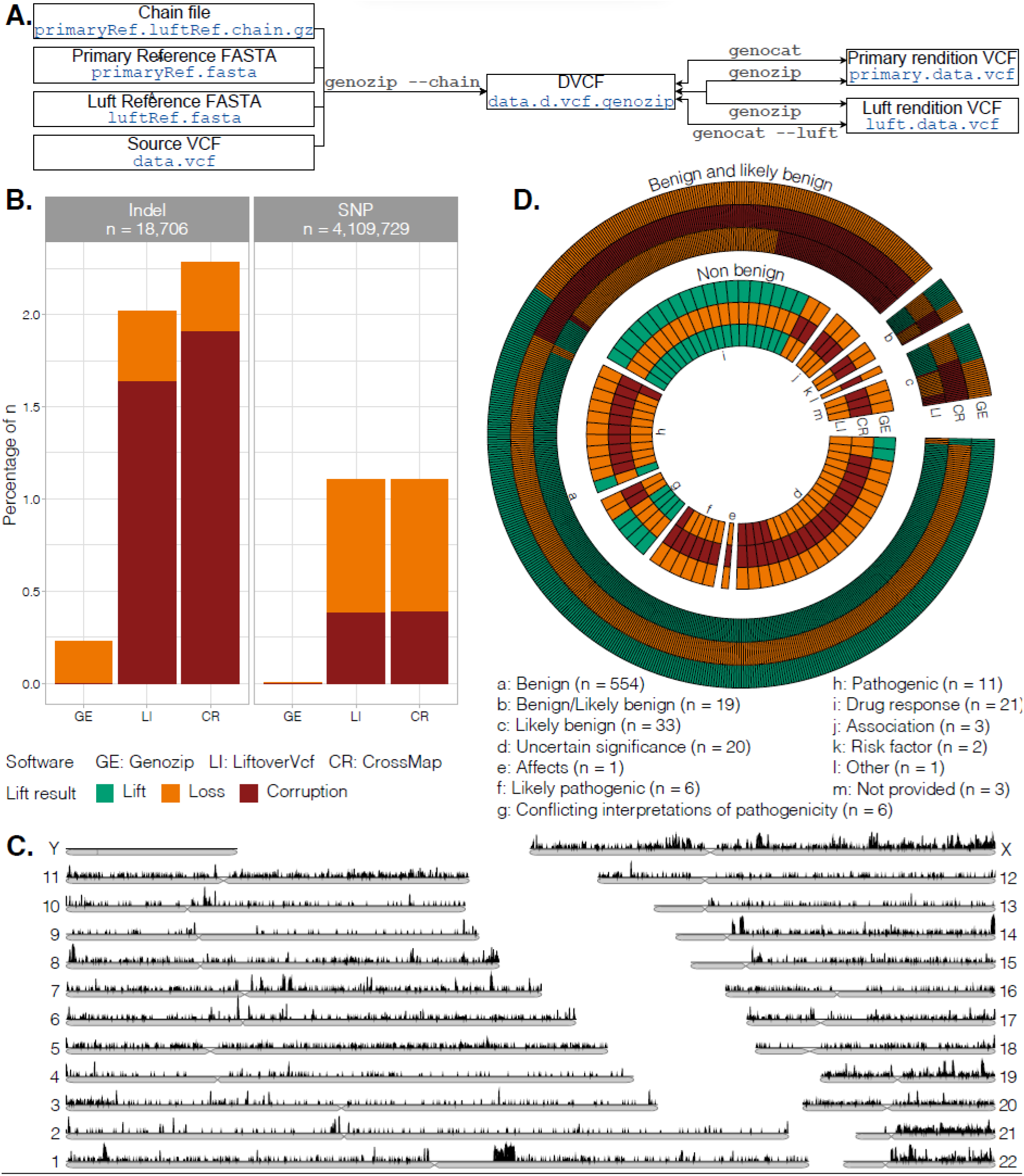
DVCF command line usage and performance vs. CrossMap and LiftoverVcf. **A**. Generation of primary (*Primary*) and alternate (*Luft*) renditions and their integration into a bioinformatics pipeline. Commands are shown along the connecting arrows and file names are indicated in blue text. File suffixes are automatically generated by genozip. **B**. When lifting ∼4.1M SNPs and ∼19k indels from GRCh37 to GRCh38 (right and left subpanels, respectively), Genozip DVCF results in substantially fewer lost variants (orange bar portions) and eradicates all forms of data corruption (red bar portions) (see Tables S1 & S9). GE: Genozip DVCF; CR: CrossMap; LI: LiftoverVcf. **C**. Genomic distribution of the ∼30k SNPs (from a set of ∼4.1M SNPs) where the reference and alternative allele are switched (i.e. REF⇄ALT allele switches) between human reference GRCh37 and GRCh38. While Genozip DVCF correctly lifted all ∼30k SNPs, CrossMap drops these variants (see Table S9), which may lead to biases in downstream analyses due to the genomic clustering of REF⇄ALT allele switches. **D**. Similar results are observed when lifting a set ∼970k variants with clinical annotations from GRCh37 to GRCh38, with CrossMap and LiftoverVcf producing many more dropped variants (orange blocks) than Genozip DVCF and also generating multiple corrupted variants (red blocks) – including pathogenic and likely pathogenic cases (see associated key) – that are entirely absent from Genozip DVCF (see tables S13-15).

While a DVCF has two possible renditions (*Primary* and *Luft*), the DVCF file format is carefully designed so that the information contained in each rendition is identical, thereby guaranteeing that the *cross-rendering* process is strictly lossless and invertible. Variants or annotations that have representation in only one of the two coordinate systems due to the lack of a chain file alignment or limitations of the lifting algorithm are nevertheless still represented in both DVCF *renditions*. The missing variants and annotations are stored in VCF header lines and certain INFO annotations, respectively, as defined in the DVCF specification (sections 5,6,7 in the specification [7]). Since the DVCF file format complies with the VCF standard, any tool that works with VCF files will also work with DVCF files. In addition to rendering the complete VCF data, Genozip’s `genocat` command allows users to render specific subsets of data in either coordinate system, as well as perform a wide variety of downsampling and filtering procedures, by drawing upon Genozip’s internal indexing facility (see the `genocat` user manual).

To compare the performance of Genozip DVCF against two widely used liftover tools, i.e., LiftoverVcf and CrossMap, we conducted a series of benchmarks using publicly available human genomic data (Supplementary Information section 2). For each file tested, we used the same chain file with all three tools, but nevertheless, each of the three tools produced a different lifted VCF file. We systematically investigated the differences in the lifted files produced by the three tools, characterising and enumerating errors and biases that result from underlying deficiencies in the algorithms of each liftover tool. In other words, we did not evaluate the correctness of the chain files, but rather the correctness of the lifting algorithms for any given chain file.

Genozip outperformed both incumbent tools when lifting a set ∼19,000 indels, or ∼4.1 million SNPs from an older version of the human reference (i.e., GRCh37) to the most recent version (i.e., GRCh38), reducing the proportion of incorrect calls by nine-fold for indels (0.2% vs 2.1–2.3%) and 463-fold for SNPs (0.002% vs 1.1%) (Fig. 1B, Tables S1-11). Notably, CrossMap also dropped all instances of SNPs where the reference and alternate alleles had been switched between the two reference versions (i.e., REF⇄ALT switches, which accounted for 0.7% of all 4.1M SNPs), which may introduce biases into downstream analyses that leverage SNP diversity patterns due to the highly non-uniform genomic distribution of the REF⇄ALT switches (Figs. 1C, S3-4). We also applied the three tools to ∼970k million clinically relevant variants obtained from the ClinVar website [10] and identified multiple cases of data corruption introduced by CrossMap and LiftoverVcf, which included known pathogenic variants, that are entirely absent from Genozip DVCF (Fig. 1D; Tables S13-15).

## Conclusion

The implementation of DVCF within the Genozip software platform provides researchers with a user-friendly and flexible tool that facilitates the construction of bioinformatic pipelines capable of working across dual coordinate systems. This is essential whenever researchers wish to exploit the advantages of working with sequence data aligned to the latest version of the reference genome, while still being able to draw upon abundant legacy data or tools that use an older reference genome version.

Genozip DVCF also correctly lifts more variants and eliminates key errors identified in our benchmarks of two widely used liftover tools, CrossMap and LiftoverVcf. Failure of these latter tools to correctly liftover variants in regions of the genome causally associated with phenotypes (Fig. 1D) could negatively impact genetic analyses that rely on regional genomic signals—such as genomic scans for disease-associated and/or selected variants. Moreover, liftover errors have documented impacts on variant effect interpretation [6], which could result in important clinically significant variants being overlooked or leading to misdiagnoses [1].

Overall, DVCF represents a new and fundamentally different approach for working concurrently within coordinate systems from two different genome assemblies—a reality that many genomics researchers will likely face as improved sequencing technologies lead to increasingly complete reference genomes [1]—greatly simplifying bioinformatic workflows without compromising the robustness of downstream analytical results. Importantly, Genozip’s extensible framework means that further improving DVCF functionality (e.g., by including new algorithms to handle liftover errors, or supporting an arbitrary number of reference genomes, thereby enabling recording of homologous genes in different species) will be an active area of future research.

## Methods

Genozip implements the Dual Coordinate VCF specification. Once a VCF file is lifted to a DVCF genozip file using the *genozip* command, that genozip file can be *rendered* into either Primary or Luft coordinates using the *genocat* command. Crucially, each one of these renditions is a DVCF file and accordingly contains all the information needed to produce both renditions—that is, these two VCF files contain identical information, just organised in a different way, and consequently can be losslessly converted back and forth. A simple example of rendering with one variant can be seen in Fig. 2. More information can be found in the Supplementary Information section 1, and in https://genozip.com/dvcf.

**Figure 2.**
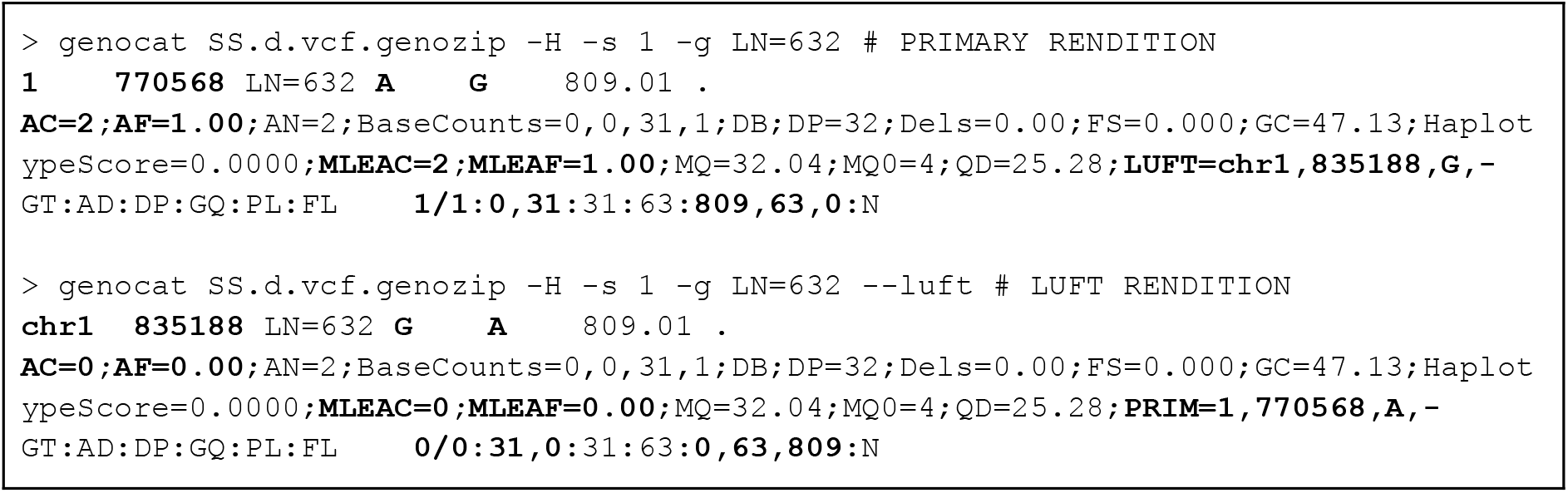
An example of rendering and cross rendering with one variant. The differences between the Primary rendition (top command line and output) and Luft rendition (bottom command line and output) are highlighted in bold font. Notice that the LUFT and PRIM subfields of the INFO field display the information that is used by Genozip to cross-render this variant.

During the lifting process, Genozip ensures that the resulting *Luft* data is correct, including the CHROM, POS, REF, and ALT fields and, crucially, all associated annotations. Genozip has built-in handling for a wide range of common annotations, with command line options also provided for the user to instruct Genozip on how to handle other annotations. Typical examples involve annotations that require their value to be changed when the REF allele switches to become the ALT allele (for example: FORMAT/GT, INFO/AF) or following a strand reversal between the two references (for example: INFO/BaseCounts). Other common occurrences include annotation changes caused by altered POS information (for example: INFO/END) and others change if the sequence quoted in REF or ALT is modified (for example: INFO/AA). In addition to making changes to the values of annotation, Genozip also transforms the names of the tags when required, for example changing between FORMAT/F1R2 and FORMAT/F2R1 due to strand reversal. As with annotation value transformations, by default Genozip handles a wide range of scenarios involving the renaming of common fields, and allows the user to define others using command line options. Finally, there are specific types of transformations not yet supported by Genozip (for example: INFO/MAX_AF). In these cases, Genozip effectively drops the affected annotation by prefixing its name with `DROP`_. A full description of how Genozip deals with annotation value changes and tag renaming can be found in https://genozip.com/dvcf-annotations.

Genozip performs coordinate lift over by first categorising each variant into one of nine types of indels or seven types of SNPs, based on the allelic states and their placement in the chain file, and applying a specific liftover method for each one of these 16 cases. For variants that cannot be classified among these 16 cases (usually a complex case, such as certain types of structural variants), Genozip adopts a conservative approach and drops the variant rather than presenting a simplistic, but likely incorrect, liftover. In Supplementary Information sections 2.2.3 through 2.2.11 we describe the nine categories of indel variants and our liftover approach for each, and contrast them with the methods we observe in LiftoverVcf and CrossMap; we do the same for the seven types of SNP variant categories in sections 2.3.3 through 2.3.9. We acknowledge that there are complex edge cases in which there could be more than one liftover outcome that might be deemed correct, and accordingly we explain the logic behind each of our choices.

## Supporting information

Genozip DVCF - Supplementary Information

## Acknowledgements

D.L. was supported by a scholarship from the University of Adelaide. Y.S. is supported by the Australian Research Council (DP190103705). R.T. is an ARC DECRA fellow (DE190101069). B.L. was an ARC Future Fellow (FT170100448).

## Contributions

DL wrote the software and the first draft of the manuscript. BL, RT and YS reviewed and edited the manuscript and provided guidance. GP provided feedback that helped improve the software.

## Supplementary Information

Supplementary Information

Supplementary text

Supplementary Figs. S1-S4

Supplementary Tables S1-S20

