## Supplementary material for "Genozip Dual-Coordinate VCF format enables efficient genomic analyses and alleviates liftover limitations": Genozip DVCF - Supplementary Information

Divon Lan[1,*], Gludhug Purnomo[1,2], Ray Tobler[1,2,3,†], Yassine Souilmi[1,4,†], Bastien Llamas[1,2,4,5,†,*]

[1] Australian Centre for Ancient DNA, School of Biological Sciences, The Environment Institute, Faculty of Sciences, The University of Adelaide, Adelaide SA 5005, Australia
[2] Centre of Excellence for Australian Biodiversity and Heritage (CABAH), School of Biological Sciences, University of Adelaide, Adelaide, SA 5005, Australia
[3] Evolution of Cultural Diversity Initiative, Australian National University, College of Asia and the Pacific, Canberra, ACT 0200, Australia
[4] National Centre for Indigenous Genomics, Australian National University, Canberra, ACT 0200, Australia
[5] Telethon Kids Institute, Adelaide, SA 5000, Australia

[†] Equal contribution

Section 1: DVCF Implementation in Genozip 2
1.1 The DVCF format 3
1.2 Genozip – a brief overview 5
1.3 The lift process 5
1.4 Rendering in Primary or Luft coordinates 6
1.5 Disk space considerations 7
Section 2: Benchmark 8
2.1 Categorizing variants by lift quality 9
2.2 Indel benchmark 9
2.2.1 Data preparation 10
2.2.2 Indel benchmark results summary 12
2.2.3 REF⇄ALT switch, entire variant mapped, no REF change 13
2.2.4 Deletion REF⇄ALT switch, REF change 17
2.2.5 Deletion REF⇄ALT switch with payload in gap 19
2.2.6 Deletion with payload partially in gap 21
2.2.7 Insertion with reverse strand 22
2.2.8 Deletion with reverse strand 25
2.2.9 REF changed but not REF⇄ALT switch 26
2.2.10 New allele when REF unchanged 27
2.2.11 New allele - not quite a REF⇄ALT switch 28
2.3 SNP benchmark 29
2.3.1 Data preparation 30
2.3.2 SNP benchmark results summary 30
2.3.3 REF⇄ALT switch in SNPs 31
2.3.4 Annotation update upon strand reversal 33
2.3.5 IUPACs 35
2.3.6 REF change, not to ALT, in bi-allelic SNPs when AF<1 36
2.3.7 REF change in multi-allelic SNPs when AF<1 37
2.3.8 REF change, not to ALT, in bi-allelic SNPs when AF=1 38
2.3.9 REF⇄ALT switch proportions 39
2.4 Benchmark summary 42
Section 3: ClinVar analysis 43
3.1 Data preparation 44
3.2 Genozip analysis 44
3.3 CrossMap analysis 45
3.4 LiftoverVcf analysis 47
3.5 ClinVar benchmark – summary 48
Section 4: GRCh38 and Telomere-to-Telomere 49
Appendix: Analysis script output 53

### Section 1: DVCF Implementation in Genozip

#### 1.1 The DVCF format

DVCF files are a type of VCF file, compliant with the VCF v4.3 specification, which contain information about variants represented in two different coordinate systems. Like any VCF file, it consists of meta-information lines prefixed with a double hash (i.e. ##), a header line prefixed with #CHROM, and data lines that each contain information about a single variant (Figure S1). Importantly, the DVCF specification[6] only defines the DVCF file format and does not prescribe liftover algorithms, making it independent of any specific software implementation (including Genozip) and we expect other bioinformatics software packages to implement DVCF as well, thereby maintaining interoperability between tools. For example, existing liftover tools could add a command line option that enables the generation of DVCF formatted outputs.

We refer to the process of converting a VCF file to the DVCF format as *lifting. Lifting* requires information external to the VCF file itself, in a format defined by the implementation. The DVCF implementation in Genozip, for example, requires two reference files—i.e. the reference file that underlies the coordinate system used in the VCF, and the alternate reference file that contains the coordinates to be lifted over—and a chain file describing the mapping of variants between these two systems, but future implementations could work differently. The coordinate system of the input VCF file is referred to as the *Primary* coordinates, and the *lifting* process updates this VCF by incorporating the required information from the alternate coordinate system, which we refer to as the *Luft* coordinate system (*Luft* being a term we introduce as an alternative past-participle of *Lift*).

Thereafter, a DVCF can be *rendered* in either *Primary* coordinates or *Luft* coordinates, and can be *cross-rendered* from *Primary* to *Luft* coordinates and vice versa. We refer to the DVCF-format VCF files rendered in the *Primary* coordinates as the *Primary rendition* and to the corresponding VCF file in the *Luft* coordinates as the *Luft rendition.* The coordinate system in which a VCF file is rendered (i.e. the coordinates represented in the CHROM and POS fields) is referred to as the *foreground* coordinate system (which may be *Primary* or *Luft*), with the non-rendered coordinate system being the *background* coordinate system.

Crucially, the information present in both *foreground* and the *background* coordinate systems is present in both renditions: in particular, the CHROM and POS fields are encoded in *foreground* coordinates and the allelic states represented in the REF and ALT fields are defined relative to the *foreground* reference genome. Simultaneously, the *background* coordinate data are encoded in the INFO/LUFT or INFO/PRIM annotation in the *Primary* or *Luft* rendition respectively (Figure S1, notes 5 and 6) for each variant and also in the meta information section (Figure S1, notes 1–5).

Annotations for some variants may differ between the *Primary* and *Luft* renditions – for example, annotations that are sensitive to a change in the reference allele (e.g. INFO/AF), or to reference file strand reversal (e.g. INFO/BaseCounts) or coordinate changes (e.g. INFO/END), will only be correct with respect to the *foreground* coordinate system. Cross-rendering of variants with rendition-specific annotations is handled by *Rendering Algorithms* (or *RendAlgs*), which are a set of algorithms that convert specific annotations between renditions to ensure consistency with the foreground coordinates. Several standard RendAlgs are defined in section 6.3 of the DVCF specification and implementation developers are encouraged, but not mandated, to use the standard RendAlgs wherever possible, but may also develop proprietary RendAlgs.

There are cases where the tag name itself, rather than the values contained in the annotation, differs between renditions—for example, the FORMAT/ADR and FORMAT/ADF tags might switch names (ADF becomes ADR and vice versa) in variants that have a reference genome strand reversal. Another example of using tag renaming would be in case dropping a particular annotation is desired. In this case, the tag name can be prefixed with DROP_. The DVCF specification defines three *Tag Renaming* attributes (see section 6.4 of the DVCF specification).

A key benefit of the DVCF file format design is that *cross-rendering* consists of simply rearranging information that is already present in the DVCF file and transforming affected annotations, and therefore this process does not require external information (such as the reference sequence or the chain files), making it computationally fast and efficient. Moreover, since DVCF files are VCF files, they can be processed in the same manner as traditional VCF files using appropriate bioinformatics tools, but provide users with the additional freedom to choose which coordinate system to use when executing specific pipeline steps without having to create duplicate VCF files, thereby reducing analytical complexity and saving considerable disk space (see section 1.5).

A likely outcome of the lifting process is that some variants may only be represented in one of the two coordinate systems due to the lack of a specific chain file alignment or liftover software limitations. To maintain information identicality across renditions, such *single-coordinate* variants are represented in both renditions, by recording their *background* coordinates in the VCF meta-information (prefixed with either `##primary_only=` or `##luft_only=` in accordance with the coordinate system that the variant is represented; see Figure S1, note 1). We refer to such variants as *Primary-only* or *Luft-only* variants depending on which reference version the variant is represented. In addition, meta information lines describing the contigs and reference file of the *background* coordinate system may also be added (DVCF specification section 5.3 and 5.6 ; Figure S1, note 3).

In conclusion, while a DVCF has two *renditions* (i.e. *Primary* and *Luft*), the DVCF file format is carefully designed so that the information contained in each *rendition* is identical, thereby guaranteeing that the *cross-rendering* between the *renditions* is a strictly lossless and invertible process.

```
[1]##primary_only=1        143163348       LN=170280       G       A       1066.01
.       AC=1;AF=0.500;AN=2;Lrej=NoMappingInChainFile   GT:AD:DP:GQ:PL:FL
0/1:152,74:1066,0,2824
[2]##contig=<ID=chr1,length=248956422>
[2]##reference=file://GRCh38_full_analysis_set_plus_decoy_hla.ref.genozip
[3]##primary_reference=file://hs37d5.ref.genozip
[3]##primary_contig=<ID=1,length=249250621>
[4]##FORMAT=<ID=AD,Number=.,Type=Integer,Description="Allelic depths for the ref
and alt alleles in the order listed",RendAlg=R>
[4]##FORMAT=<ID=GT,Number=1,Type=String,Description="Genotype",RendAlg=GT>
[4]##FORMAT=<ID=PL,Number=G,Type=Integer,Description="Normalized, Phred-scaled
likelihoods for genotypes as defined in the VCF specification",RendAlg=G>
[4]##INFO=<ID=AC,Number=A,Type=Integer,Description="Allele count in genotypes, for
each ALT allele, in the same order as listed",RendAlg=A_AN>
[4]##INFO=<ID=AF,Number=A,Type=Float,Description="Allele Frequency, for each ALT
allele, in the same order as listed",RendAlg=A_1>
[4]##INFO=<ID=AN,Number=1,Type=Integer,Description="Total number of alleles in
called genotypes",RendAlg=NONE>
[5]##INFO=<ID=LUFT,Number=4,Type=String,Description="Info for rendering variant in
LUFT coords. See
https://genozip.com/dvcf.html",Source="genozip",Version="12.0.8",RendAlg=NONE>
[5]##INFO=<ID=PRIM,Number=4,Type=String,Description="Info for rendering variant in
PRIMARY coords",Source="genozip",Version="12.0.-1",RendAlg=NONE>
[5]##INFO=<ID=Lrej,Number=1,Type=String,Description="Reason variant was rejected
for LUFT coords",Source="genozip",Version="12.0.-1",RendAlg=NONE>
[5]##INFO=<ID=Prej,Number=1,Type=String,Description="Reason variant was rejected
for PRIMARY coords",Source="genozip",Version="12.0.-1",RendAlg=NONE>
#CHROM  POS     ID      REF     ALT     QUAL    FILTER  INFO    FORMAT  SS6004478
[6]chr1    597733  LN=298  A       G       263.03  .
AC=2;AF=1.00;AN=2;PRIM=1,533113,A,-  GT:AD:PL       1/1:4,20:263,24,0
```

**Figure S1**. A small subset of the Luft rendition of our SNP test file (see section 4), containing two variants. Notes: 1a Primary-only variant represented in a ##primary_only line. 2foreground (Luft in this case) coordinate data 3background (Primary) meta-information lines are preserved with the prefix primary_. 4INFO and FORMAT meta-information lines contain the RendAlg attribute 5Four DVCF INFO tags defined. 6PRIM contains the data needed to cross-render to Primary.

#### 1.2 Genozip – a brief overview

Genozip is a software platform that stores genomic files in a compressed format. For VCF files, compression is typically 2–8 times better than other standard compression tools such as gzip and BCF (Lan *et al.*, 2020). In Genozip's implementation of DVCF files, data are stored in the Genozip format. Users may render the data in either coordinate system and have the option of piping the data through an analytical tool or pipeline before returning the data to Genozip format (see Figure 1A). In the following section, we outline the basic commands involved in the *lifting* process and provide a brief description of the algorithms involved in *cross-rendering* between coordinate systems.

#### 1.3 The *lift* process

Genozip can lift one or more VCF files into a DVCF using a suitable chain file available from Ensembl (https://ftp.ensembl.org/pub/assembly_mapping/) or the UCSC Genome Browser (https://hgdownload.soe.ucsc.edu/downloads.html) by invoking the following command:

```
genozip --chain mychainfile.chain.genozip myvariants.vcf
```

This will generate a DVCF file in Genozip format: myvariants.d.vcf.genozip.

Genozip accepts chain files in the UCSC chain format (genome.ucsc.edu/goldenPath/help/chain.html), which must first be compressed with `genozip` using Primary and Luft reference files:

```
genozip --reference primary.ref.genozip --reference
luft.ref.genozip mychainfile.chain
```

Each reference file, in turn, is produced from the relevant reference genome FASTA file:
`genozip --make-reference primary.fa.gz`

During the lift process (i.e. executing `genozip --chain`), Genozip determines whether it is possible to represent each variant in both coordinate systems, with all variants failing this process (i.e. rejected variants) being retained as Primary-only variants. A variant may be rejected for three reasons: (1) it lacks an alignment in the chain file, or (2) the lifted variant has more than two alleles (either because it had more than two alleles in the input VCF file, or the Luft REF allele is neither the Primary REF nor ALT allele), or (3) if any of the INFO or FORMAT annotations cannot be cross-rendered due to the "Rejected If" condition of their RendAlg (see next section). Known cases in which Genozip cannot lift variants are listed in genozip.com/dvcf-limitations.html.

#### 1.4 Rendering in *Primary* or *Luft* coordinates

When in the Genozip format, DVCF files may be rendered in Primary or Luft coordinates by using either `genocat data.vcf.genozip` or `genocat --luft data.vcf.genozip`, respectively (Figure S2). The resulting VCF file is sorted according to its foreground (i.e., rendered) coordinates. Running `genozip` on the rendered file in either coordinate system produces a compressed DVCF file in Genozip format containing identical information.

In addition to variants becoming single coordinate during the initial lift process, they can also become single coordinate when compressing a DVCF rendition with the `genozip` command if (1) new variants were added to the VCF following the initial lifting step and these variants lack the INFO/PRIM or INFO/LUFT fields (i.e. they are single-coordinate), or (2) if an annotation was added or modified in a way that satisfies its RendAlg's "Rejected If" condition (see next paragraph).

Since both renditions are DVCF files, each rendition contains all the information needed for rendering in either coordinate system. Cross-rendering involves the translation of annotations between the Primary and Luft coordinate systems. This process often implicates making rendition-specific changes to one or more annotations for some variants, with each annotation being handled by a Rendering Algorithm (or RendAlg). RendAlgs are stand-alone algorithms that are not tied a priori to specific INFO and FORMAT tags, rather, they are assigned to tags using a new RendAlg attribute added to each ##INFO and ##FORMAT meta-information line. Thereafter, all INFO and FORMAT annotations (in the VCF data lines) of a specific tag will be treated with the `RendAlg` assigned to that tag. Importantly, Genozip adds the `RendAlg` attribute to each tag's meta-information line based on its ID and/or Number attributes, but only if the `RendAlg` attribute is not already present. This allows users to assign RendAlgs differently than Genozip's built-in assignments.

Genozip implements eleven RendAlgs, listed in genozip.com/dvcf-rendering.html, which are an implementation of the DVCF standard set of RendAlgs defined in the specification6section 6.3; see also genozip.com/dvcf-rendering.html). Each RendAlg is comprised of three elements: (1) a trigger; which is the class of event handled by the RendAlg, (2) an action which is the effect the RendAlg has on the annotation, and (3) "Rejected If" conditions, which describe circumstances in which the designated action cannot be applied despite the trigger activating, in which case the variant is rejected (i.e., is represented as a single-coordinate variant in the DVCF). To illustrate this mechanism, consider the A_1 RendAlg which is useful for annotations in fields reporting allele frequency information. The A_1 RendAlg trigger is a "REF⇄ALT switch" (i.e., a change of REF allele across renditions), its action is to recalculate the value of the annotation as 1 - value, and the variant is "Rejected If" the value of the annotation is outside of the range [0,1] or if the variant has more than two alleles. By default, Genozip assigns the A_1 RendAlg to the `FORMAT/AF`, `INFO/AF`, `INFO/MLEAF`, `INFO/LDAF` fields and any additional INFO tag that begins with `AF_` or ends with `_AF` (except `MAX_AF`). Another example is the RendAlg named END that Genozip assigns by default to the `INFO/END` field. Its trigger event is "Always" (i.e., it triggers on all `INFO/END` annotations in the data), its action is to modify the annotation to which it is assigned to maintain the same distance from POS in both renditions, whereby the variant is "Rejected If" the position indicated by the annotation and POS are not both on the same chain file alignment. A final example is the XREV RendAlg whose trigger event is "strand reversal" (as indicated by the chain file alignment) and its action is to reverse the elements of an array. Genozip assigns the XREV RendAlg to the `INFO/BaseCounts` field.

```
> genocat SS.d.vcf.genozip -H -s 1 -g LN=632   # PRIMARY RENDITION
1       770568  LN=632  A       G       809.01   .
AC=2;AF=1.00;AN=2;BaseCounts=0,0,31,1;DB;DP=32;Dels=0.00;FS=0.000;GC=47.1
3;HaplotypeScore=0.0000;MLEAC=2;MLEAF=1.00;MQ=32.04;MQ0=4;QD=25.28;LUFT=c
hr1,835188,G,-  GT:AD:DP:GQ:PL:FL       1/1:0,31:31:63:809,63,0:N

> genocat SS.d.vcf.genozip -H -s 1 -g LN=632 --luft # LUFT RENDITION
chr1    835188  LN=632  G       A       809.01   .
AC=0;AF=0.00;AN=2;BaseCounts=0,0,31,1;DB;DP=32;Dels=0.00;FS=0.000;GC=47.1
3;HaplotypeScore=0.0000;MLEAC=0;MLEAF=0.00;MQ=32.04;MQ0=4;QD=25.28;PRIM=1
,770568,A,-     GT:AD:DP:GQ:PL:FL       0/0:31,0:31:63:0,63,809:N
```

**Figure S2**. An example of rendering and cross rendering with one variant. The differences between the Primary rendition (top command line and output) and Luft rendition (bottom command line and output) are highlighted in bold font. Notice that the LUFT and PRIM subfields of the INFO field display the information that is used by Genozip to cross-render this variant.

#### 1.5 Disk space considerations

In addition to needing only a single DVCF file to represent variants in two coordinate systems, rather than two separate VCF files, Genozip also stores VCF files using highly efficient compression (Lan et al. 2020). Together, this amounts to significant saving of disk space.

To demonstrate this, we compare the size of our SNP and Indel test files (SI section 1):

**Table S16:** SNP file

| | **Uncompressed** | **.gz compressed** | **Genozip DVCF** |
|---|---|---|---|
| GRCh37[1] | 999 MB | 199 MB | 76 MB |
| GRCh38[2] | 1094 MB | 175 MB | |
| **Total** | **2093 MB** | **374 MB** | **76 MB** |
| **Compression ratio** | **-** | **5.6X** | **27.5X** |

[1]SS6004478.annotated.nh2.variants.vcf [2]snp.38.gatk.vcf

**Table S17:** Indel file

| | **Uncompressed** | **.gz compressed** | **Genozip DVCF** |
|---|---|---|---|
| GRCh37[1] | 3961 KB | 848 KB | 443 KB |
| GRCh38[2] | 4470 KB | 777 KB | |
| **Total** | **8431 KB** | **1625 KB** | **443 KB** |
| **Compression ratio** | | **5.2X** | **19X** |

[1]indel.37.vcf [2]indel.38.gatk.vcf

### Section 2: Benchmark

#### 2.1 Categorizing variants by lift quality

We used chromosome 22 data from the 1000 Genome project phase 1 for benchmarking indels (see Section 1.2) and a sample from https://sharehost.hms.harvard.edu/genetics/reich_lab/sgdp/vcf_variants/ for benchmarking SNPs (see Section 1.3).

We categorize the 18,857 indel variants in the indel test file and the 4,109,729 variants in the SNP test file, in respect to *lifting*—the operation of converting the VCF file from one coordinate system to another, GRCh37 to GRCh38 in our case.

With respect to each of the three tools, Genozip, LiftoverVcf and CrossMap, we assign one of five categories to each variant. Numbers 1 and 2 are good outcomes, and numbers 3 through 5 are bad outcomes, with increasing order of severity.

1) ***Lifted*** - The lift operation succeeded and the resulting variant is correct.

2) ***Unmapped*** - The chain file has no mapping for the coordinate of this variant, and therefore the variant was correctly rejected from lifting by the tool.

3) ***Annotation Loss*** - The variant was lifted and the resulting variant is correct but incomplete—some of the annotations it originally contained were dropped by the tool.

4) ***Variant Loss*** - The variant was rejected from lifting by the tool, despite having all the information needed to successfully lift it. This happens when variants have some complexities that are beyond the capabilities of the particular tool.

5) ***Data Corruption*** - The variant was lifted, but the resulting variant contains incorrect data.

In some cases, we decided to categorize a variant as *Lifted* even though it had a very *Minor data loss*. This was done in cases where data loss almost certainly will not affect any downstream analysis. These cases are explained where they occur in the sections below.

#### 2.2 Indel benchmark

##### 2.2.1 Data preparation

We developed a set of scripts that executes this benchmark in its entirely - from downloading the data to producing the analysis files. It is available from https://github.com/divonlan/genozip-dvcf-results and the entry point script is: run-indl-37-38.sh

The key steps executed by this script are these:

***Preparing the input files***

The indel test file was generated from the chromosome 22 VCF file of the 1000 Genome Project phase 1 obtained from ftp://ftp-trace.ncbi.nih.gov/1000genomes/ftp/release/20110521/ALL.chr22.phase1_release_v3.20101123.snps_indels_svs.genotypes.vcf.gz and filtered to contain only its indel variants (N=18,706 indels).

The GRCh37 reference was obtained from ftp://ftp.1000genomes.ebi.ac.uk/vol1/ftp/technical/reference/phase2_reference_assembly_sequence/hs37d5.fa.gz, and prepared for Genozip use with

```
> genozip --make-reference hs37d5.fa.gz
```

The GRCh38 reference was downloaded from ftp://ftp.1000genomes.ebi.ac.uk/vol1/ftp/technical/reference/GRCh38_reference_genome/GRCh38_full_analysis_set_plus_decoy_hla.fa, and prepared for Genozip use with:

```
> genozip --make-reference GRCh38_full_analysis_set_plus_decoy_hla.fa.gz -o
GRCh38.ref.genozip
```

The chain file mapping GRCh37 genomic features to their corresponding GRCh38 coordinates was obtained from http://ftp.ensembl.org/pub/assembly_mapping/homo_sapiens/GRCh37_to_GRCh38.chain.gz, and prepared with:

```
> genozip --echo GRCh37_to_GRCh38.chain.gz --reference hs37d5.ref.genozip
--reference GRCh38.ref.genozip --match-chrom-to-reference --force --output
GRCh37_to_GRCh38.matched.chain.genozip
```

Note the `--match-chrom-to-reference`: this modifies the contig names in the chain file to match those of the reference files (e.g. “22” vs “chr22”).

For our indels-test, we used a single sample, indels-only VCF file that was generated with the following commands:

```
> genozip
ALL.chr22.phase1_release_v3.20101123.snps_indels_svs.genotypes.vcf.gz
```

```
> genocat
ALL.chr22.phase1_release_v3.20101123.snps_indels_svs.genotypes.vcf.genozip
--samples 1 --indels-only -o indel.37.vcf
```

***Genozip DVCF liftover***

Using the chain file, which itself uses the two reference files, we generated a DVCF file. For the purpose of ease of comparison between corresponding VCF files in the two coordinate systems, we replaced the ID field with a line number (e.g. "LN=452") using the `--add-line-numbers` command line option, and modified the contig names to match those of the reference file and the chain file with `--match-chrom-to-reference`.

```
> genozip --echo --chain
GRCh37_to_GRCh38.matched.chain.genozip --add-line-numbers
--match-chrom-to-reference indel.37.vcf -o
indel-37-38/indel-37-38.d.vcf.genozip
```

In order to test the other tools with the VCF file that contains the updated ID field and contig names, and also to allow is comparison of Genozip DVCF status to the outcome of the other tools, we generated a GRCh37-only version of the VCF that contains the DVCF lifting status (which we refer to as "oStatus"). The `--single` option converts the dual-coordinate DVCF file to a single-coordinate (GRCh37 in this case) VCF file. `--show-ostatus` adds an `INFO/oSTATUS` field to each variant, describing the Genozip's liftover status of this variant. The ID field already contains the line number.

```
> genocat indel-37-38.d.vcf.genozip --single -o
indel.37.annotated.vcf --show-ostatus
```

***CrossMap liftover***

CrossMap.py version v0.5.2 was installed using conda and the test VCF file was lifted over to GRCh38 coordinates using the following command:

```
> CrossMap.py vcf GRCh37_to_GRCh38.matched.chain
indel-37-38/indel.37.annotated.vcf
GRCh38_full_analysis_set_plus_decoy_hla.fa.gz
indel-37-38/indel.38.CrossMap.vcf
```

***GATK LiftoverVcf liftover***

GATK version 4.1.7 was used, and the indel test file lifted over to GRCh38 coordinates was produced using `LiftoverVcf`:

```
> gatk --java-options '-Xmx16g -XX:ParallelGCThreads=1'
LiftoverVcf --INPUT indel-37-38/indel.37.annotated.vcf
--OUTPUT indel-37-38/indel.38.gatk.vcf --CHAIN
GRCh37_to_GRCh38.matched.chain --REJECT
indel-37-38/indel.38.gatk.rejects.vcf
--RECOVER_SWAPPED_REF_ALT --REFERENCE_SEQUENCE
GRCh38_full_analysis_set_plus_decoy_hla.fa.gz
--TAGS_TO_REVERSE AF --TAGS_TO_REVERSE AF_EUR
--TAGS_TO_REVERSE AF_EAS --TAGS_TO_REVERSE AF_AMR
--TAGS_TO_REVERSE AF_SAS --TAGS_TO_REVERSE AF_AFR
--TAGS_TO_REVERSE AF_EUR_unrel --TAGS_TO_REVERSE AF_EAS_unrel
--TAGS_TO_REVERSE AF_AMR_unrel --TAGS_TO_REVERSE AF_SAS_unrel
--TAGS_TO_REVERSE AF_AFR_unrel --TAGS_TO_DROP AC
--TAGS_TO_DROP AC_EUR --TAGS_TO_DROP AC_EAS --TAGS_TO_DROP
AC_AMR --TAGS_TO_DROP AC_SAS --TAGS_TO_DROP AC_AFR
--TAGS_TO_DROP AC_EUR_unrel --TAGS_TO_DROP AC_EAS_unrel
--TAGS_TO_DROP AC_AMR_unrel --TAGS_TO_DROP AC_SAS_unrel
--TAGS_TO_DROP AC_AFR_unrel --TAGS_TO_DROP AC_Hom_EUR
--TAGS_TO_DROP AC_Hom_EAS --TAGS_TO_DROP AC_Hom_AMR
--TAGS_TO_DROP AC_Hom_SAS --TAGS_TO_DROP AC_Hom_AFR
--TAGS_TO_DROP AC_Hom --TAGS_TO_DROP AC_Het_EUR --TAGS_TO_DROP
AC_Het_EAS --TAGS_TO_DROP AC_Het_AMR --TAGS_TO_DROP AC_Het_SAS
--TAGS_TO_DROP AC_Het_AFR --TAGS_TO_DROP AC_Het --TAGS_TO_DROP
AC_Hom_EUR_unrel --TAGS_TO_DROP AC_Hom_EAS_unrel
--TAGS_TO_DROP AC_Hom_AMR_unrel --TAGS_TO_DROP
AC_Hom_SAS_unrel --TAGS_TO_DROP AC_Hom_AFR_unrel
--TAGS_TO_DROP AC_Het_EUR_unrel --TAGS_TO_DROP
AC_Het_EAS_unrel --TAGS_TO_DROP AC_Het_AMR_unrel
--TAGS_TO_DROP AC_Het_SAS_unrel --TAGS_TO_DROP
AC_Het_AFR_unrel
```

Finally, our bash scripts perform analyses and output three summary files, one for each tool (`analysis.CrossMap.txt`, `analysis.gatk.txt`, `analysis.Genozip.txt`), which describe the outcome of the variants (Lifted or Dropped) categorised from Genozip's oStatus categories. See Supplementary Information section 6 for the content of these files.

#### 2.2.2 Indel benchmark results summary

**Table S1**. Performance of three liftover tools for indels, including handling of problematic variants, according to standard categories reported by the Genozip software.

| # Variants | Category | Genozip | LiftoverVcf | CrossMap |
|---|---|---|---|---|
| 18,201 | Lifted without issues | ✓*Lifted* | ✓*Lifted* | ✓*Lifted* |
| 78 | REF not mapped in chain file | ✓*Unmapped* | ✓*Unmapped* | ✓*Unmapped* |
| 153 | REF⇄ALT switch but REF unchanged (switch in number of repeats) | ✓*Lifted* | ✗ *Data Corruption* | ✗ *Data Corruption* |
| 27 | REF⇄ALT switch but REF unchanged (Flanking regions indicate switch) | ✓*Lifted* | ✗ *Data Corruption* | ✗ *Data Corruption* |
| 6 | Simple Deletion->Insertion REF⇄ALT switch | ✓*Lifted* | ✗ *Variant Loss* | ✗ *Data corruption* |
| 71 | Deletion->Insertion REF⇄ALT switch called by Deletion payload in chain file gap | ✓*Lifted* | ✗ *Variant Loss* | ✗ *Variant Loss* |
| 20 | Deletion with payload partially in gap | ✓*Unmapped* | ✓*Unmapped* | ✗ *Data Corruption* |
| 40 | Insertion with reverse strand | ✓*Lifted* | ✗ *Data Corruption (35)* ✓*Lifted (5)* | ✗ *Data Corruption* |
| 67 | Deletion with reverse strand | ✓*Lifted* | ✗ *Data Corruption (51)* ✓*Lifted (16)* | ✗ *Data Corruption* |
| 9 | Apparent REF<>ALT switch disqualified by flanking regions | ✗ *Variant Loss* | ✗ *Data Corruption* | ✗ *Data Corruption* |
| 13 | REF changed in Deletion but not REF⇄ALT switch | ✗ *Variant Loss* | ✗ *Variant Loss* | ✗ *Data Corruption* |
| 11 | REF unchanged, but flanking regions show that this is a new Insertion allele. | ✗ *Variant Loss* | ✗ *Data Corruption* | ✗ *Data Corruption* |
| 6 | The REF base mismatches between the references | ✗ *Variant Loss* | ✗ *Variant Loss* | ✗ *Data Corruption* |
| 4 | REF unchanged, but flanking regions show that this is a new Deletion allele. | ✗ *Variant Loss* | ✗ *Data Corruption* | ✗ *Data Corruption* |
| **18,706** | **TOTAL** | | | |

##### 2.2.3 REF⇄ALT switch, entire variant mapped, no REF change

From Table S1:

| # Variants | Genozip | LiftoverVcf | CrossMap |
|---|---|---|---|
| 180 | ***Lifted*** | ***Data Corruption*** | ***Data Corruption*** |

**Genozip oSTATUS:** `OkRefAltSwitchIndelRpts`, `OkRefAltSwitchIndelFlank`

We found that both LiftoverVcf and CrossMap fail to identify 180 REF⇄ALT switches in indel variants, respectively, resulting in incorrect variants with the REF and ALT reversed vs their correct values, along with errors in many of the INFO and FORMAT annotations that depend on the order of alleles (such as AC, AF, AD, GL, etc.). Genozip lifts all 186 REF⇄ALT switches,

**Case 1**: simple insertion with REF⇄ALT switch

Consider this GRCh37 insertion variant, which maps to 16423118 in GRCh38:

```
#CHROM   POS        ID       REF      ALT
22       16903845   LN=133   T        TAC
```

```
> genocat --reference hs37d5.ref.genozip -r chr22:16903845+8
16903845-16903852          TAGAGGCA
```

```
> genocat --reference GRCh38.ref.genozip -r chr22:16423118+10
16423118-16423127          TACAGAGGCA
```

It is easy to see that the insertion got incorporated in the GRCh38 reference, therefore the haplotypes with GT=1 in the original VCF, indicating their samples have this insertion, are the REF allele in the Luft reference, while those haplotypes that don't have this insertion would be a deletion variant relative to the Luft reference. In other words, this is a REF⇄ALT switch.

Genozip correctly executes a REF⇄ALT switch (Table S2; note the fields that differ from the original VCF in red), while both LiftoverVcf and CrossMap fail to detect this REF⇄ALT switch and therefore generate a data-corrupted variant.

**Table S2:** Example VCF of an insertion with REF⇄ALT switch.

| | Genozip (37) | Genozip (38) | LiftoverVcf (38) | CrossMap (38) |
|---|---|---|---|---|
| | genocat indel-37-38.d.vcf.genozip --luft --samples 1 --no-header -g LN=133 | genocat indel-37-38.d.vcf.genozip --luft --samples 1 --no-header -g LN=133 | grep -w LN=133 indel.38.gatk.vcf \|cut -f1-10 | grep -w LN=133 indel.38.CrossMap.vcf \|cut -f1-10 |
| #CHROM | 22 | **chr22** | **chr22** | 22 |
| POS | 16903845 | **16423118** | **16423118** | **16423118** |
| ID | LN=133 | LN=133 | LN=133 | LN=133 |
| REF | T | **TAC** | T | T |
| ALT | TAC | **T** | TAC | TAC |
| QUAL | 166 | 166 | 166 | 166 |
| FILTER | PASS | PASS | PASS | PASS |
| INFO | AA=TAC;ERATE=0.0118;RSQ=0.5469;AN=2184;LDAF=0.2750;VT=INDEL;THETA=0.0057;AVGPOST=0.7502;AC=455;AF=0.21;ASN_AF=0.14;AMR_AF=0.19;AFR_AF=0.44;EUR_AF=0.12;**LUFT=chr22,16423118,TAC,-** | AA=TAC;ERATE=0.0118;RSQ=0.5469;AN=2184;**LDAF=0.7250**;VT=INDEL;THETA=0.0057;AVGPOST=0.7502;**AC=1729**;**AF=0.79;ASN_AF=0.86;AMR_AF=0.81;AFR_AF=0.56;EUR_AF=0.88;PRIM=22,16903845,T,-** | AA=TAC;AC=455;AF=0.21;AFR_AF=0.44;AMR_AF=0.19;AN=2184;ASN_AF=0.14;AVGPOST=0.7502;ERATE=0.0118;EUR_AF=0.12;LDAF=0.2750;RSQ=0.5469;THETA=0.0057;VT=INDEL | AA=TAC;ERATE=0.0118;RSQ=0.5469;AN=2184;LDAF=0.2750;VT=INDEL;THETA=0.0057;AVGPOST=0.7502;AC=455;AF=0.21;ASN_AF=0.14;AMR_AF=0.19;AFR_AF=0.44;EUR_AF=0.12 |
| FORMAT | GT:DS:GL | GT:DS:GL | GT:DS:GL | GT:DS:GL |
| HG00096 | 0\|0:0.050:0.00,-1.20,-18.40 | **1\|1:1.950:-18.40,-1.20,0.00** | 0\|0:0.050:0.00,-1.20,-18.40 | 0\|0:0.050:0.00,-1.20,-18.40 |

GRCh version is in parentheses in the headers. The VCF lines are presented transposed, for readability. **Red**: changes vs. the original VCF; highlight: errors.

**Case 2:** Insertion with repeats with REF⇄ALT switch

Consider the variant at POS=22780917 that maps to 22426580 on GRCh38:

```
#CHROM  POS       ID       REF    ALT
22      22780917  LN=3114  C      CA
```

Viewing the Primary (37) and Luft references: Notice that the Insertion was incorporated in the Luft reference—it has 3 As instead of 2:

```
> genocat -e hs37d5.ref.genozip -r chr22:22780917+9
22780917-22780925        CAATCGGTC

> genocat -e GRCh38.ref.genozip -r chr22:22426580+10
22426580-22426589        CAAATCGGTC
```

Genozip executed a REF⇄ALT switch while the other tools failed to do so (Table S3).

**Table S3:** Example VCF of an insertion with repeats with REF⇄ALT switch.

| | **Genozip (37)** | **Genozip (38)** | **LiftoverVcf (38)** | **CrossMap (38)** |
|---|---|---|---|---|
| | genocat indel-37-38.d.vcf.genozip -H -s1 -g LN=3114 | genocat indel-37-38.d.vcf.genozip -H -s1 --luft -g LN=3114 | grep -w LN=3114 indel.38.gatk.vcf \|cut -f1-10 | grep -w LN=3114 indel.38.CrossMap.vcf \|cut -f1-10 |
| #CHROM | 22 | **chr22** | **chr22** | 22 |
| POS | 22780917 | **22426580** | **22426580** | **22426580** |
| ID | LN=3114 | LN=3114 | LN=3114 | LN=3114 |
| REF | C | **CA** | C | C |
| ALT | CA | **C** | CA | CA |
| QUAL | 763 | 763 | 763 | 763 |
| FILTER | PASS | PASS | PASS | PASS |
| INFO | AA=.;ERATE=0.0005;AN=2184;VT=INDEL;THETA=0.0005;AC=1517;LDAF=0.6930;RSQ=0.9505;AVGPOST=0.9681;AF=0.69;ASN_AF=0.63;AMR_AF=0.56;AFR_AF=0.86;EUR_AF=0.71;**LUFT=chr22,22426580,CA,-** | AA=.;ERATE=0.0005;AN=2184;VT=INDEL;THETA=0.0005;**AC=667;LDAF=0.3070**;RSQ=0.9505;AVGPOST=0.9681;**AF=0.31;ASN_AF=0.37;AMR_AF=0.44;AFR_AF=0.14;EUR_AF=0.29;PRIM=22,22780917,C,-** | AA=.;AC=1517;AF=0.69;AFR_AF=0.86;AMR_AF=0.56;AN=2184;ASN_AF=0.63;AVGPOST=0.9681;ERATE=0.0005;EUR_AF=0.71;LDAF=0.6930;RSQ=0.9505;THETA=0.0005;VT=INDEL | AA=.;ERATE=0.0005;AN=2184;VT=INDEL;THETA=0.0005;AC=1517;LDAF=0.6930;RSQ=0.9505;AVGPOST=0.9681;AF=0.69;ASN_AF=0.63;AMR_AF=0.56;AFR_AF=0.86;EUR_AF=0.71 |
| FORMAT | GT:DS:GL | GT:DS:GL | GT:DS:GL | GT:DS:GL |
| HG00096 | 0\|1:1.000:-7.40,0.00,-6.20 | **1\|0:1.000:-6.20,0.00,-7.40** | 0\|1:1.000:-7.40,0.00,-6.20 | 0\|1:1.000:-7.40,0.00,-6.20 |

GRCh version is in parentheses in the headers. The VCF lines are presented transposed, for readability. **Red**: changes vs. the original VCF; highlight: errors.

**Case 3:** Deletion with repeats with REF⇄ALT switch

Consider the variant at POS=24483878 that maps to 24087925 on GRCh38:

```
#CHROM   POS        ID        REF      ALT
22       24483878   LN=4150   AT       A
```

Viewing the Primary (37) and Luft (38) references: Notice that the Deletion was incorporated in the Luft reference - it has only 2 repeating Ts instead of 3:

```
> genocat -e hs37d5.ref.genozip -r chr22:24483878+10
24483878-24483887         ATTTAGGGAC

> genocat -e GRCh38.ref.genozip -r chr22:24087925+9
24087925-24087933         ATTAGGGAC
```

Genozip executed a REF⇄ALT switch while the other tools failed to do so (Table S4).

**Table S4:** Example VCF of a deletion with REF⇄ALT switch.

| | **Genozip (37)** | **Genozip (38)** | **LiftoverVcf (38)** | **CrossMap (38)** |
|---|---|---|---|---|
| | genocat indel-37-38.d.vcf .genozip -H -s 1 -g LN=4150 | genocat indel-37-38.d.vcf .genozip -H -s 1 --luft -g LN=4150 | grep -w LN=4150 indel.38.gatk.vcf \| cut -f1-10 | grep -w LN=4150 indel.38.CrossMap. vcf \| cut -f1-10 |
| #CHROM | 22 | chr22 | chr22 | 22 |
| POS | 24483878 | 24087925 | 24087925 | 24087925 |
| ID | LN=4150 | LN=4150 | LN=4150 | LN=4150 |
| REF | AT | A | AT | AT |
| ALT | A | AT | A | A |
| QUAL | 785 | 785 | 785 | 785 |
| FILTER | PASS | PASS | PASS | PASS |
| INFO | AA=AT;AC=1441;AN=2184;LDAF=0.6585;VT=INDEL;THETA=0.0006;AVGPOST=0.9913;RSQ=0.9854;ERATE=0.0006;AF=0.66;ASN_AF=0.55;AMR_AF=0.82;AFR_AF=0.46;EUR_AF=0.80;LUFT=chr22,24087925,A,- | AA=AT;AC=743;AN=2184;LDAF=0.3415;VT=INDEL;THETA=0.0006;AVGPOST=0.9913;RSQ=0.9854;ERATE=0.0006;AF=0.34;ASN_AF=0.45;AMR_AF=0.18;AFR_AF=0.54;EUR_AF=0.20;PRIM=22,24483878,AT,- | AA=AT;AC=1441;AF=0.66;AFR_AF=0.46;AMR_AF=0.82;AN=2184;ASN_AF=0.55;AVGPOST=0.9913;ERATE=0.0006;EUR_AF=0.80;LDAF=0.6585;RSQ=0.9854;THETA=0.0006;VT=INDEL | AA=AT;AC=1441;AN=2184;LDAF=0.6585;VT=INDEL;THETA=0.0006;AVGPOST=0.9913;RSQ=0.9854;ERATE=0.0006;AF=0.66;ASN_AF=0.55;AMR_AF=0.82;AFR_AF=0.46;EUR_AF=0.80 |
| FORMAT | GT:DS:GL | GT:DS:GL | GT:DS:GL | GT:DS:GL |
| HG00096 | 0\|1:1.000:-2.20,0.00,-15.80 | 1\|0:1.000:-15.80,0.00,-2.20 | 0\|1:1.000:-2.20,0.00,-15.80 | 0\|1:1.000:-2.20,0.00,-15.80 |

GRCh version is in parentheses in the headers. The VCF lines are presented transposed, for readability. Red: changes vs. the original VCF; highlight: errors.

##### 2.2.4 Deletion REF⇄ALT switch, REF change

**From Table S1:**

| # Variants | Genozip | LiftoverVcf | CrossMap |
|---|---|---|---|
| 6 | ***Lifted*** | ***Variant Loss*** | ***Data Corruption*** |

**Genozip oSTATUS:** `OkRefAltSwitchDelToIns`

We found that 6 variants with REF⇄ALT switch resulting in a REF change are either lost (LiftoverVcf) or mishandled leading to Data Corruption (CrossMap). Genozip successfully lifted these variants.

Consider the variant at POS=17998325 that maps to 17519295 on GRCh38:

```
#CHROM   POS        ID         REF      ALT
22       17998325   LN=825     TTG      T
```

Viewing the Primary (37) and Luft (38) references in the local context:

```
> genocat -e hs37d5.ref.genozip -r chr22:17998325+10
17998322-17998344        GTTTTGTTTTTTTTTTTTGAGAC

> genocat -e GRCh38.ref.genozip -r chr22:17519295 +10
17519292-17519314        GTTTTTTTTTTTTTTTTTGAGAC
```

The variant observed in the samples might be a true deletion or might indeed mirror the variation between the references, in which case it would have been better categorized as a SNP rather than a deletion. Regardless, given that this is categorized as a deletion in the VCF on hand, Genozip declares it a REF⇄ALT switch since the haplotypes with this variant become the REF allele in the Luft reference, and the ones without the variant are the ALT allele.

In cases like this where the REF changes between the references (here: TTG->TTT), LiftoverVcf rejects the variant with “MismatchedRefAllele”, whereas CrossMap simply updates the REF to TTT causing a Data Corruption (because the REF allele no longer represents the true sequences of the haplotypes with GT=0) (Table S5).

**Table S5:** Example VCF of a REF⇄ALT switch with REF change.

| | **Genozip (37)** | **Genozip (38)** | **LiftoverVcf (38)** | **CrossMap (38)** |
|---|---|---|---|---|
| | genocat indel-37-38.d.vcf .genozip -H -s 1 -g LN=825 | genocat indel-37-38.d.vcf .genozip -H -s 1 --luft -g LN=825 | Variant lost | grep -w LN=825 indel.38.CrossMap.vcf \| cut -f1-10 |
| #CHROM | 22 | **chr22** | | 22 |
| POS | 17998325 | **17519295** | | **17519295** |
| ID | LN=825 | LN=825 | | LN=825 |
| REF | TTG | **T** | | **TTT** |
| ALT | T | **TTG** | | T |
| QUAL | 191 | 191 | | 191 |
| FILTER | PASS | PASS | | PASS |
| INFO | AA=.;AC=1970;AF=0.90;AFR_AF=0.99;AMR_AF=0.85;AN=2184;ASN_AF=0.96;AVGPOST=0.9826;ERATE=0.0014;EUR_AF=0.83;LDAF=0.8963;RSQ=0.9295;THETA=0.0003;VT=INDEL;LUFT=chr22,17519295,T,- | AA=.;**AC=214;AF=0.10;AFR_AF=0.01;AMR_AF=0.15**;AN=2184;**ASN_AF=0.04**;AVGPOST=0.9826;ERATE=0.0014;**EUR_AF=0.17;LDAF=0.1037**;RSQ=0.9295;THETA=0.0003;VT=INDEL**;PRIM=22,17998325,TTG,-** | | AA=.;AC=1970;AF=0.90;AFR_AF=0.99;AMR_AF=0.85;AN=2184;ASN_AF=0.96;AVGPOST=0.9826;ERATE=0.0014;EUR_AF=0.83;LDAF=0.8963;RSQ=0.9295;THETA=0.0003;VT=INDEL |
| FORMAT | GT:DS:GL | GT:DS:GL | | GT:DS:GL |
| HG00096 | 1\|1:2.000:-9.00,-1.70,0.00 | **0\|0:0.000:0.00,-1.70,-9.00** | | 1\|1:2.000:-9.00,-1.70,0.00 |

GRCh version is in parentheses in the headers. The VCF lines are presented transposed, for readability. **Red**: changes vs. the original VCF; highlight: errors.

##### 2.2.5 Deletion REF⇄ALT switch with payload in gap

**From Table S1:**

| # Variants | Genozip | LiftoverVcf | CrossMap |
|---|---|---|---|
| 71 | *Lifted* | *Variant Loss* | *Variant Loss* |

**Genozip oSTATUS:** `OkRefAltSwitchWithGap`

When a deletion variant in the Primary reference enters the Luft reference, in other words, the deletion payload that existed in the Primary reference no longer exists in the Luft reference, this will often manifest itself as a gap in the chain file. We found 71 of these variants; lifting over with LiftoverVcf and CrossMap resulted in Variant Loss, while Genozip handled these variants correctly.

Consider for example:

```
#CHROM   POS        ID          REF          ALT
22       17995661   LN=821      TTTGCTGTTG   T
```

In the chain file we have the following alignments:

```
> genocat GRCh37_to_GRCh38.chain.genozip --show-chain
…
Primary: 22 17995313-17995661 Luft: chr22 17516283-17516631 Xstrand=-
Primary: 22 17995671-17996285 Luft: chr22 17516632-17517246 Xstrand=-
…
```

As can be appreciated, the anchor base of the variant, **T**, is the final base on the first alignment (that ends at 17995661), and the entire 9-base payload, **TTGCTGTTG**, precisely fits in the gap between the alignments (17995662 to 17995670).

When inspecting the two references starting at the anchor base **T**, it is clear that the Luft reference incorporates this deletion, and therefore the variant in a REF⇄ALT switch:

```
> genocat --reference hs37d5.ref.genozip -r chr22:17995661+15
17995661-17995675         TTTGCTGTTGTTGCC

> genocat -reference GRCh38.ref.genozip -r chr22:17516631+6
17516631-17516636         TTTGCC
```

These variants are correctly categorized as a REF⇄ALT switch by Genozip. However, LiftoverVcf rejects them with “NoTarget” and CrossMap rejects them with “Fail(REF==ALT)”, leading to Variant Loss (Table S6).

**Table S6:** Example VCF of a REF⇄ALT switch with payload in gap.

| | Genozip (37) | Genozip (38) | LiftoverVcf (38) | CrossMap (38) |
|---|---|---|---|---|
| | genocat indel-37-38.d.vcf.genozip -r 17995661 -s HG00104 --header-one | genocat indel-37-38.d.vcf.genozip -r 17516631 -s HG00104 --luft | Variant lost | Variant lost |
| #CHROM | 22 | chr22 | | |
| POS | 17995661 | 17516631 | | |
| ID | LN=821 | LN=821 | | |
| REF | TTTGCTGTTG | T | | |
| ALT | T | TTTGCTGTTG | | |
| QUAL | 1144 | 1144 | | |
| FILTER | PASS | PASS | | |
| INFO | AA=TTTGCTGTTG;ERATE=0.0124;AN=2184;VT=INDEL;THETA=0.0005;AVGPOST=0.9345;AC=1914;LDAF=0.8546;RSQ=0.8066;AF=0.88;ASN_AF=0.94;AMR_AF=0.84;AFR_AF=0.95;EUR_AF=0.80;LUFT=chr22,17516631,T,- | AA=TTTGCTGTTG;ERATE=0.0124;AN=2184;VT=INDEL;THETA=0.0005;AVGPOST=0.9345;AC=270;LDAF=0.1454;RSQ=0.8066;AF=0.12;ASN_AF=0.06;AMR_AF=0.16;AFR_AF=0.05;EUR_AF=0.20;PRIM=22,17995661,TTTGCTGTTG,- | | |
| FORMAT | GT:DS:GL | GT:DS:GL | | |
| HG00104 | 0\|1:0.550:0.00,-1.20,-41.70 | 1\|0:1.450:-41.70,-1.20,0.00 | | |

GRCh version is in parentheses in the headers. The VCF lines are presented transposed, for readability. **Red**: changes vs. the original VCF.

#### 2.2.6 Deletion with payload partially in gap

**From Table S1:**

| # Variants | Genozip | LiftoverVcf | CrossMap |
|---|---|---|---|
| 20 | ***Variant Loss*** | ***Variant Loss*** | ***Data Corruption*** |

**Genozip oSTATUS:** `REFSplitInChain`

When a deletion payload is partially in a chain file gap (other than the case where the entire payload is in the gap), CrossMap updates the REF by removing the bases that fall in the gap. This is incorrect: while with the lifted REF the haplotypes that contain the variant (GT=1) are now correctly represented, the haplotypes with GT=0 are incorrectly represented as the actual sequenced data contains the previous REF, not the lifted one. Rather, this should be a new allele.

In contrast, since neither Genozip nor LiftoverVcf are capable of adding an allele they both correctly reject these variants (with the resulting Variant Loss).

Example: the indel test file:
```
#CHROM   POS        ID          REF       ALT
22       17995306   LN=818      ATTATAT   A
```

CrossMap-lifted variant:
```
#CHROM   POS        ID          REF       ALT
22       17516277   LN=818      ATTATA    A
```

The chain file has the last base in REF (POS=17995312) in the gap between two alignments:
```
Primary: chr22 17992953-17995311 Luft: chr22 17513924-17516282
Primary: chr22 17995313-17995661 Luft: chr22 17516283-17516631
```

#### 2.2.7 Insertion with reverse strand

**From Table S1:**

| # Variants | Genozip | LiftoverVcf | CrossMap |
|---|---|---|---|
| 40 | ***Lifted*** | ***Data Corruption (35)***<br>***Lifted (5)*** | ***Data Corruption*** |

**Genozip oSTATUS:** `OkRefSameInsRev`

Consider the following Insertion variant in our indel test file (in GRCh37 coordinates):

```
#CHROM   POS        ID     REF    ALT     INFO(partial)
22       16566319   LN=75  A      ACAAT   AC=13;AF=0.01;AN=2184
```

Per the chain file, POS maps to 15411644 in GRCh38, on the reverse strand. CrossMap lifts this insertion by reverse complementing the REF and ALT:

```
#CHROM   POS        ID     REF    ALT     INFO(partial)
chr22    15411644   LN=75  T      ATTGT   AC=13;AF=0.01;AN=2184
```

While the CrossMap-lifted variant contains precisely the same information as the original variant, it is unfortunately non-compliant with the VCF specification (section 5.2) that requires the variant's anchor base (**A** lifted to **T**) be on the left side, whereas here it is right-anchored. This is likely to break downstream tools.

LiftoverVcf goes further, and left-aligns the resulting variant, leading to this:

```
#CHROM   POS        ID     REF    ALT     INFO(partial)
chr22    15411637   LN=75  C      CTGAT   AC=13;AF=0.01;AN=2184
```

The algorithm LiftoverVcf applied here is as follows:
GRCh38 (Luft) in region 15411637-15411644 is **CTGATTGT**. Since **TGATTG** is 1½ repeats of the insertion variant payload **ATTG** (traversing backwards from the anchor base **T**), it seems that we can represent this variant in a canonical left-aligned way with **CTGAT** at POS=15411637, because the original insertion without left-aligning **CTGATTG**$^{\mathbf{ATTG}}$**T** yields precisely the same sequence as the insertion after left aligning: **C**$^{\mathbf{TGAT}}$**TGATTGT**.

However, this is wrong. The reason is that there is no requirement in the VCF specification that a VCF file must contain all the variants of its specific samples, and it is not true that any loci lacking a variant is an indication that all the samples in the VCF file have a base equal to the reference at that locus. Only the loci covered by the variants listed in the VCF file are known, and we cannot make any assumption regarding the bases the specific samples in the VCF file have at other loci.

Consider, for example, the LiftoverVcf-generated left-aligned variant above. This variant now asserts that the 13 haplotypes (AC=13) with GT=1 for this variant, have the bases **CTGAT** starting at position 15411637. This is simply not knowable from the data at hand, and might, in fact, be wrong for any of the 13 haplotypes. For these 13 haplotypes in this

VCF file, we know nothing at all about their bases in loci 15411637–15411643, all we know is that they have an insertion of **ATTG** just before locus 15411644.

This is therefore a risk of Data Corruption: if any one of the 13 haplotypes doesn't have **CTGAT** bases starting at 15311637, or if any one of the other 2171 haplotypes doesn't have a **C** at this locus, then the VCF file is corrupted.

In addition to the above issue of Data Corruption, there is an additional issue of Data Loss: the original VCF informed us that all 2184 haplotypes have a **T** at 15411644, and that 13 haplotypes have a **ATTG** just before 15411644. This information is no longer present in the LiftoverVcf-generated file, and therefore lost.

Genozip chooses to left-anchor the variant, but not left-align it:

| #CHROM | POS | ID | REF | ALT | INFO(partial) |
|---|---|---|---|---|---|
| chr22 | 15411643 | LN=75 | G | **GATTG** | AC=13;AF=0.01;AN=2184 |

This is quite similar to the CrossMap variant, except Genozip's anchor base **G** is to the left, rather than the right, of the insertion payload **ATTG**, which is also reflected in POS. This makes it compliant with the VCF specification, while not risking Data Corruption.

Note that this solution is still not perfect: namely, it asserts that all samples in the VCF have **G** at 15411643, which is not known from the data, and it loses the information that all samples have a **T** at 15411644. However, without access to the full sequence of all samples, or alternatively knowledge that the VCF contains all variants in the samples (and taking into account neighboring variants in the computation), this is the best that can be done. We consider this issue to be a *Minor data loss* that does not affect the categorization.

All 40 variants with reverse strand Insertion were affected for CrossMap, but only 33 of them for LiftoverVcf - the remaining 7 were cases where left-aligning resulted in the same variant as left-anchoring.

An additional Data Corruption issue is that if an INFO/AA field exists, both CrossMap and LiftoverVcf fail to update it appropriately. This issue affects 14 of the 40 variants, including 2 of the 7 for which LiftoverVcf correctly left-anchored. See the example in Table S7.

Finally, we note a *Minor Data Loss* in LiftoverVcf is due to conversion of the FORMAT/GL field to FORMAT/PL, with the loss of granularity. As discussed, a *Minor Data Loss* does not affect the categorization.

**Table S7:** Example VCF of insertion with reverse strand.

| | Genozip (37) | Genozip (38) | LiftoverVcf (38) | CrossMap (38) |
|---|---|---|---|---|
| | genocat indel-37-38.d.vcf .genozip -H -s 1 -g LN=107 | genocat indel-37-38.d.vcf. genozip -H -s 1 --luft -g LN=107 | grep -w LN=107 indel.38.gatk.vcf \| cut -f1-10 | grep -w LN=107 indel.38.CrossMap .vcf \| cut -f1-10 |
| #CHROM | 22 | **chr22** | **chr22** | 22 |
| POS | 16687501 | **15290461** | **15290461** | **15290462** |
| ID | LN=107 | LN=107 | LN=107 | LN=107 |
| REF | C | **C** | **C** | **G** |
| ALT | CA | **CT** | **CT** | **TG** |
| QUAL | 208 | 208 | 208 | 208 |
| FILTER | PASS | PASS | PASS | PASS |
| INFO | AA=CA;AC=103;AF=0.05;AFR_AF=0.07;AMR_AF=0.04;AN=2184;ASN_AF=0.05;AVGPOST=0.9793;ERATE=0.0006;EUR_AF=0.03;LDAF=0.0530;RSQ=0.8577;THETA=0.0152;VT=INDEL;**LUFT=chr22,15290461,C,X** | **AA=CT**;AC=103;AF=0.05;AFR_AF=0.07;AMR_AF=0.04;AN=2184;ASN_AF=0.05;AVGPOST=0.9793;ERATE=0.0006;EUR_AF=0.03;LDAF=0.0530;RSQ=0.8577;THETA=0.0152;VT=INDEL;**PRIM=22,16687501,C,X** | AA=CA;AC=103;AF=0.05;AFR_AF=0.07;AMR_AF=0.04;AN=2184;ASN_AF=0.05;AVGPOST=0.9793;ERATE=0.0006;EUR_AF=0.03;LDAF=0.0530;RSQ=0.8577;**ReverseComplementedAlleles**;THETA=0.0152;VT=INDEL | AA=CA;AC=103;AF=0.05;AFR_AF=0.07;AMR_AF=0.04;AN=2184;ASN_AF=0.05;AVGPOST=0.9793;ERATE=0.0006;EUR_AF=0.03;LDAF=0.0530;RSQ=0.8577;THETA=0.0152;VT=INDEL |
| FORMAT | GT:DS:GL | GT:DS:GL | GT:DS:**PL** | GT:DS:GL |
| HG00096 | 0\|0:0.000:0.00,-0.60,-8.40 | 0\|0:0.000:0.00,-0.60,-8.40 | 0\|0:0.000:**0,6,84** | 0\|0:0.000:0.00,-0.60,-8.40 |
| | | | 0\|1:1.050:48,3,0 | 0\|1:1.050:-4.80,-0.30,0.00 |

GRCh version is in parentheses in the headers. The VCF lines are presented transposed, for readability. **Red**: changes vs. the original VCF; highlight: errors.

#### 2.2.8 Deletion with reverse strand

**From Table S1:**

| # Variants | Genozip | LiftoverVcf | CrossMap |
|---|---|---|---|
| 67 | *Lifted* | *Data Corruption (51)*<br>*Lifted (16)* | *Data Corruption* |

**Genozip oSTATUS:** `OkRefSameDelRev`

This issue is similar to the previous one, just for deletions rather than insertions.

Consider the following deletion, in GRCh37 coordinates:

```
#CHROM   POS         ID      REF     ALT    INFO(partial)
22       16524572    LN=70   ACACT   A      AC=112;AF=0.05;AN=2184
```

CrossMap lifts it by reverse-complementing REF and ALT and moving POS from what became REF's right-most base, 15453391, to its left-most base 15453387:

```
#CHROM   POS         ID      REF     ALT    INFO(partial)
chr22    15453387    LN=70   AGTGT   T      AC=112;AF=0.05;AN=2184
```

As discussed for insertions, this variant contains correct information, however it is non-compliant with the VCF specification, and hence we will consider it a Data Corruption.

LiftoverVcf, as in insertions, goes further, and left-aligns the resulting variant, creating the following variant:

```
#CHROM  POS         ID      REF     ALT    INFO(partial)
chr22   15453384   LN=70   CTGAG   C      AC=112;AF=0.05;AN=2184
```

Brief explanation: GRCh38, chr22, region 15453384-15453391 is: **CTGAGTGT**. The deletion generated by the original (CrossMap) variant **CTG**$_{\text{AGTG}}$**T** results in an identical sequence as the deletion described by the LiftoverVcf's left-aligned variant: **C**$_{\text{TGAG}}$**TGT**. However, as before, this is wrong because it makes possibly incorrect assumptions about the nucleotide sequences of the samples at loci 15453384–15453386 which are not in fact knowable from the data.

Again, similar to the insertion case, Genozip left-anchors but does not left-align the variant, which is the optimal (yet still imperfect) solution:

```
#CHROM   POS         ID      REF     ALT    INFO(partial)
chr22    15453386    LN=70   GAGTG   G      AC=112;AF=0.05;AN=2184
```

All 67 variants with reverse strand deletion were affected for CrossMap, but only 46 of them for LiftoverVcf—the remaining 21 were cases where left-aligning resulted in the same variant as left-anchoring, however, 5 of the 21 are nevertheless corrupted due to failure to update the INFO/AA field, bringing the total of corrupted LiftoverVcf variants to 51.

##### 2.2.9 REF changed but not REF⇄ALT switch

**From Table S1:**

| # Variants | Genozip | LiftoverVcf | CrossMap |
|---|---|---|---|
| 19 | ***Variant Loss*** | ***Variant Loss*** | ***Data Corruption*** |

**Genozip oSTATUS:** `RefNewAlleleDelRefChanged`, `RefNewAllelInsRefChanged`

In indels, if the bases of the REF change between the two references, CrossMap simply updates the REF field with the new bases. This is completely wrong, and is responsible for corruption of 26 variants in our test file. In contrast, both Genozip and LiftoverVcf reject the variants in this case (losing their data).

**Example 1 (Insertion)**:

Consider the following insertion variant in our indel test file (in GRCh37 coordinates):

```
#CHROM   POS        ID       REF      ALT      INFO(partial)
22       18068419   LN=871   A        ATT      AC=311;AF=0.14;AN=2184
```

POS=18068419 in GRCh37 maps to 17585653 in GRCh38, and this position has a base change—from A in GRCh37 to T in GRCh38. Because of this base change, both Genozip and LiftoverVcf reject this variant. However, CrossMap produced the following:

```
#CHROM   POS        ID       REF      ALT      INFO(partial)
22       17585653   LN=871   T        ATT      AC=311;AF=0.14;AN=2184
```

This is obviously completely wrong. First, it changed a left-anchored insertion (with an A anchor base) to a right-anchored insertion (with a T anchor base). Second, it is asserting that all 311 haplotypes (AC=311) with GT=1 have a T at this position, while in fact the original VCF informs us that they have an A. Finally, it is an invalid insertion variant per the VCF specification 5.2.

**Example 2 (Deletion)**:

Consider the following deletion variant in our indel test file (in GRCh37 coordinates):

```
#CHROM   POS        ID       REF      ALT      INFO(partial)
22       18068421   LN=872   TA       T        AC=631;AF=0.29;AN=2184
```

The CrossMap lifted variant:

```
#CHROM   POS        ID       REF      ALT      INFO(partial)
22       17585655   LN=872   TT       T        AC=631;AF=0.29;AN=2184
```

The reference has a base change in the second base of REF. Therefore, this would be a new allele and the correct variant would be REF=TT ALT=TA,T. Since neither Genozip nor LiftoverVcf are capable of adding an allele, they both reject this variant. However, CrossMap's variant asserts that all the (AN-AC)=1553 haplotypes with GT=0 have a TT at this position, while in fact they have a TA.

##### 2.2.10 New allele when REF unchanged

**From Table S1:**

| # Variants | Genozip | LiftoverVcf | CrossMap |
|---|---|---|---|
| 15 | *Variant Loss* | *Data Corruption* | *Data Corruption* |

**Genozip oSTATUS:** `RefNewAlleleInsSameRef`, `RefNewAlleleDelSameRef`

Cases where the bases of REF are identical in both references, yet the new reference contains a new allele. These variants are lifted incorrectly by both CrossMap and LiftoverVcf yielding corrupted variants. In contrast, Genozip rejects them because it is not capable of adding an allele.

Example:

Original variant:

```
#CHROM   POS        ID          REF      ALT
22       22735735   LN=3091     T        TG
```

CrossMap and LiftoverVcf variant in GRCh38—unchanged REF and ALT:

```
#CHROM   POS        ID          REF      ALT
chr22    22381366   LN=3091     T        TG
```

Looking at the references:

```
> genocat -e hs37d5.ref.genozip -r chr22:22735735+10
22735735-22735744       TAGGGAACTG

> genocat -e GRCh38.ref.genozip -r chr22:22381366+10
22381366-22381375       TGGGGAACTG
```

At first glance, this might look like a REF⇄ALT switch since TG is present in the Luft reference. However, looking at the variant in its local context, it is clear that the Luft reference represents a new allele that is neither REF nor ALT, and the new variant would be:

```
#CHROM   POS        ID       REF      ALT
chr22    22381366   LN=3091  TG       TA,TAG
```

Genozip correctly identifies this as a case of a new allele, and rejects the variant as it is not capable of adding another allele. CrossMap and LiftoverVcf in contrast, incorrectly lift the variant resulting in Data Corruption.

#### 2.2.11 New allele - not quite a REF⇄ALT switch

**From Table S1:**

| # Variants | Genozip | LiftoverVcf | CrossMap |
|---|---|---|---|
| 9 | *Variant Loss* | *Data Corruption* | *Data Corruption* |

**Genozip oSTATUS:** `RefNewAlleleIndelNoSwitch`

Consider the following case:

```
#CHROM  POS       ID       REF     ALT
22      22484247  LN=2870  A       AC
```

Inspecting both references, with 4 flanking bases, we see:

Primary reference: CAGG**A**AATG

Luft reference: CAGG**AC**AGTG

As first glance, it appears to be a REF⇄ALT switch where the insertion AC got incorporated in the Luft reference. However, Genozip also compares the flanking 4 bases on either side to verify that is indeed a REF⇄ALT switch. In this case, the region to the right is different - an AATG in the Primary reference vs AGTG in the Luft reference, and therefore Genozip rejects the REF⇄ALT switch hypothesis and instead determines that the Luft reference represents a new allele that is neither the REF nor the ALT. Since Genozip cannot currently add new alleles, it rejects this variant with `RefNewAlleleIndelNoSwitch`.

We also experimented with a value of 2 for the length of the flanking regions to be tested. When this more permissive approach is used, we observed the following changes in the oSTATUS categories of the variants in the indel test file (Table S8).

**Table S8:** Variant categorisation changes if changing the flanking regions test from four bases on either side, to two.

| oSTATUS | 4 bases | 2 bases |
|---|---|---|
| RefNewAlleleIndelNoSwitch | 9 | 2 |
| OkRefAltSwitchIndelFlank | 27 | 34 |
| RefNewAlleleDelRefChanged | 13 | 10 |
| RefNewAlleleInsSameRef | 17 | 11 |
| OkRefAltSwitchDelToIns | 6 | 9 |
| RefNewAllelInsRefChanged | 0 | 6 |
| Other categories - no change | | |

#### 2.3 SNP benchmark

##### 2.3.1 Data preparation

We developed a set of scripts that executes this benchmark in its entirely - from downloading the data to producing the analysis files. It is available from https://github.com/divonlan/genozip-dvcf-results and the entry point script is: run-snp-37-38.sh

The key steps executed by this script are these:

The SNP test file, SS6004478.annotated.nh2.variants.vcf.gz, containing 4,109,729 SNP variants, was extracted from the tar archive: https://sharehost.hms.harvard.edu/genetics/reich_lab/sgdp/vcf_variants/vcfs.variants.public_samples.279samples.tar.

We used the same reference files and chain file as the indel test.

Similar to the indel test, we proceed to preparing the DVCF:

```
> genozip --chain GRCh37_to_GRCh38.chain.genozip
SS6004478.annotated.nh2.variants.vcf.gz --add-line-numbers
--match-chrom-to-reference -o snp-37-38.d.vcf.genozip
```

We then proceed to generate a GRCh37-only file with contains the add lines numbers, the updated contig names and adding the INFO/oSTATUS field to each variant, reporting Genozip's lift-over status:

```
> genocat snp-37-38.d.vcf.genozip --single --show-ostatus -o
snp.37.annotated.vcf
```

Testing CrossMap: As in the indel test, CrossMap.py version v0.5.2 was used:

```
> CrossMap.py vcf GRCh37_to_GRCh38.chain.gz snp.37.annotated.vcf
GRCh38_full_analysis_set_plus_decoy_hla.fa snp.38.CrossMap.vcf
```

Testing GATK LiftoverVCF: As in the indel test, LiftoverVcf contained in GATK version 4.17 was used to lift over the snp test file to GRCh38:

```
> gatk --java-options '-Xmx16g -XX:ParallelGCThreads=1' LiftoverVcf
--INPUT snp.37.annotated.vcf --OUTPUT snp.38.gatk.vcf --CHAIN
GRCh37_to_GRCh38.matched.chain --REJECT snp.38.gatk.rejects.vcf
--RECOVER_SWAPPED_REF_ALT --REFERENCE_SEQUENCE
GRCh38_full_analysis_set_plus_decoy_hla.fa.gz --TAGS_TO_REVERSE AF
--TAGS_TO_REVERSE MLEAF --TAGS_TO_DROP AC --TAGS_TO_DROP MLEAC
```

As in the indel test, our bash script produces three files: `analysis.CrossMap.txt` `analysis.gatk.txt` `analysis.Genozip.txt`.

#### 2.3.2 SNP benchmark results summary

**Table S9**. Performance of three liftover tools for SNPs, including handling of problematic variants, according to standard categories reported by the Genozip software. Note that LiftoverVcf contains command line options that determine the handling of REF⇄ALT switches. Variants can either be i) dropped (the default, which result in Variant Loss), ii) kept, with updated annotations for a subset of variants that LiftoverVcf is able to revise (resulting in Data Corruption due to a subset of variants maintaining incorrect annotations), or iii) kept, with some annotations being converted or dropped (resulting in Annotation Loss). We chose the latter option in our benchmarks, which we consider to be the least problematic.

| # Vars. | Category | Genozip | LiftoverVcf | CrossMap |
|---|---|---|---|---|
| 4,037,520 | No issues - lifted | ✓ ***Lifted*** | ✓ ***Lifted*** | ✓ ***Lifted*** |
| 26,728 | No issues - no mapping | ✓ ***Unmapped*** | ✓ ***Unmapped*** | ✓ ***Unmapped*** |
| 29,635 | REF⇄ALT switch in SNPs | ✓ ***Lifted*** | ✗ ***Annotation Loss*** | ✗ ***Variant Loss*** |
| 15,689 | Annotation update upon strand reversal | ✓ ***Lifted*** | ✗ ***Data Corruption*** | ✗ ***Data Corruption*** |
| 12 | IUPACs | ✓ ***Lifted*** | ✗ ***Variant Loss*** | ✗ ***Data Corruption*** |
| 68 | REF change, not to ALT, in bi-allelic SNPs when AF<1 | ✗ ***Variant Loss*** | ✗ ***Variant Loss*** | ✗ ***Data Corruption*** |
| 30 | REF change in multi-allelic SNPs when AF<1 | ✗ ***Variant Loss*** | ✗ ***Variant Loss*** | ✗ ***Data Corruption*** |
| 47 | REF change, not to ALT, in bi-allelic SNPs when AF=1 | ✓ ***Lifted*** | ✗ ***Variant Loss*** | ✓ ***Lifted*** |
| **4,109,729** | **TOTAL** | | | |

##### 2.3.3 REF⇄ALT switch in SNPs

**From Table S9:**

| # Variants | Genozip | LiftoverVcf | CrossMap |
|---|---|---|---|
| 29635 | ***Lifted*** | ***Annotation Loss*** | ***Variant Loss*** |

**Genozip oSTATUS:** `OkRefAltSwitchSNP`

In case of a REF⇄ALT switch in a bi-allelic SNP, Genozip lifts the variant, updating all the relevant annotations. CrossMap drops all these variants. LiftoverVcf drops these variants by default, but is capable of lifting them with the --RECOVER_SWAPPED_REF_ALT option, however this it offers very limited corrections (with --TAGS_TO_REVERSE and --TAGS_TO_DROP), which would cause data loss (if fields are dropped) or corruption (if they are not).

In the following example (Table S10), we used LiftoverVcf options
`--RECOVER_SWAPPED_REF_ALT --TAGS_TO_REVERSE AF`
`--TAGS_TO_REVERSE MLEAF`.

LiftoverVcf correctly updates the fields INFO/AF, INFO/MLEAF and FORMAT/GT, FORMAT/AD, FORMAT/PL but fails to correct INFO/AC, INFO/MLEAC. We can avoid a Data Corruption due to AC and MLEAC by also applying --TAGS_TO_DROP AC and --TAGS_TO_DROP MLEAC, and hence we categorize LiftoverVcf as Annotation Loss for these variants.

We also note that LiftoverVcf has an additional *Minor Data Loss* due to re-ordering of the FORMAT annotations, thereby losing the information of their original order.

**Table S10:** Example VCF of REF⇄ALT switch in a SNP.

| | Genozip (37) | Genozip (38) | LiftoverVcf (38) | CrossMap (38) |
|---|---|---|---|---|
| | genocat snp-37-38.d.vcf.genozip -H -s 1 -g LN=632 | genocat snp-37-38.d.vcf.genozip -H -s 1 --luft -g LN=632 | grep -w LN=632 snp.38.gatk.vcf \| cut -f1-10 | |
| #CHROM | 1 | **chr1** | **chr1** | |
| POS | 770568 | **835188** | **835188** | |
| ID | LN=632 | LN=632 | LN=632 | |
| REF | A | **G** | **G** | |
| ALT | G | **A** | **A** | |
| QUAL | 809.01 | 809.01 | 809.01 | |
| FILTER | . | . | **PASS** | |
| INFO | AC=2;AF=1.00;AN=2;BaseCounts=0,0,31,1;DB;DP=32;Dels=0.00;FS=0.000;GC=47.13;HaplotypeScore=0.0000;MLEAC=2;MLEAF=1.00;MQ=32.04;MQ0=4;QD=25.28;**LUFT=chr1,835188,G,-** | **AC=0;AF=0.00**;AN=2;BaseCounts=0,0,31,1;DB;DP=32;Dels=0.00;FS=0.000;GC=47.13;HaplotypeScore=0.0000;**MLEAC=0;MLEAF=0.00**;MQ=32.04;MQ0=4;QD=25.28;**PRIM=1,770568,A,-** | **AF=0.00**;AN=2;BaseCounts=0,0,31,1;DB;DP=32;Dels=0.00;FS=0.000;GC=47.13;HaplotypeScore=0.0000;**MLEAF=0.00**;MQ=32.04;MQ0=4;QD=25.28;**SwappedAlleles** | |
| FORMAT | GT:AD:DP:GQ:PL:FL | GT:AD:DP:GQ:PL:FL | GT:AD:DP:FL:GQ:PL | |
| SS6004478 | 1/1:0,31:31:63:809,63,0:N | **0/0:31,0**:31:63:**0,63,809**:N | **0/0:31,0**:31:N:63:**0,63,809** | |

GRCh version is in parentheses in the headers. The VCF lines are presented transposed, for readability. **Red**: changes vs. the original VCF; highlight: errors.

##### 2.3.4 Annotation update upon strand reversal

**From Table S9:**

| # Variants | Genozip | LiftoverVcf | CrossMap |
|---|---|---|---|
| 15689 | ***Lifted*** | ***Data Corruption*** | ***Data Corruption*** |

**Genozip oSTATUS:** `OkRefSameSNPRev`

Sometimes when REF, ALT are reverse-complemented due to the chain file mapping being to the reverse strand, it is necessary to update some annotations. Both LiftoverVcf and CrossMap fail to do so, resulting in data-corrupted variants. In our test file, the affected annotation is INFO/BaseCounts annotation (Table S11).

**Table S11:** Example VCF of annotation update upon strand reversal.

| | **Genozip (37)** | **Genozip (38)** | **LiftoverVcf (38)** | **CrossMap (38)** |
|---|---|---|---|---|
| | genocat snp-37-38.d.vcf.genozip -H -s 1 -g LN=253 | genocat snp-37-38.d.vcf.genozip -H -s 1 --luft -g LN=253 | grep -w LN=253 snp.38.gatk.vcf \| cut -f1-10 | grep -w LN=253 snp.38.CrossMap.vcf \| cut -f1-10 |
| #CHROM | 1 | **chr1** | **chr1** | 1 |
| POS | 364127 | **455210** | **455210** | **455210** |
| ID | LN=253 | LN=253 | LN=253 | LN=253 |
| REF | G | **C** | **C** | **C** |
| ALT | A | **T** | **T** | **T** |
| QUAL | 26.78 | 26.78 | 26.78 | 26.78 |
| FILTER | . | . | PASS | . |
| INFO | AC=2;AF=1.00;AN=2;BaseCounts=5,0,23,0;BaseQRankSum=0.804;DB;DP=28;Dels=0.00;FS=0.000;GC=38.65;HaplotypeScore=0.0000;MLEAC=2;MLEAF=1.00;MQ=4.61;MQ0=26;MQRankSum=0.804;QD=0.96;ReadPosRankSum=0.804;**LUFT=chr1,455210,C,X** | AC=2;AF=1.00;AN=2;**BaseCounts=0,23,0,5**;BaseQRankSum=0.804;DB;DP=28;Dels=0.00;FS=0.000;GC=38.65;HaplotypeScore=0.0000;MLEAC=2;MLEAF=1.00;MQ=4.61;MQ0=26;MQRankSum=0.804;QD=0.96;ReadPosRankSum=0.804;**PRIM=1,364127,G,X** | AC=2;AF=1.00;AN=2;BaseCounts=5,0,23,0;BaseQRankSum=0.804;DB;DP=28;Dels=0.00;FS=0.000;GC=38.65;HaplotypeScore=0.0000;MLEAC=2;MLEAF=1.00;MQ=4.61;MQ0=26;MQRankSum=0.804;QD=0.96;ReadPosRankSum=0.804;**ReverseComplementedAlleles** | AC=2;AF=1.00;AN=2;BaseCounts=5,0,23,0;BaseQRankSum=0.804;DB;DP=28;Dels=0.00;FS=0.000;GC=38.65;HaplotypeScore=0.0000;MLEAC=2;MLEAF=1.00;MQ=4.61;MQ0=26;MQRankSum=0.804;QD=0.96;ReadPosRankSum=0.804 |
| FORMAT | GT:AD:DP:GQ:PL:FL | GT:AD:DP:GQ:PL:FL | GT:AD:DP:FL:GQ:PL | GT:AD:DP:GQ:PL:FL |
| SS6004478 | 1/1:23,5:27:3:25,3,0:N | 1/1:23,5:27:3:25,3,0:N | 1/1:23,5:27:N:3:25,3,0 | 1/1:23,5:27:3:25,3,0:N |

GRCh version is in parentheses in the headers. The VCF lines are presented transposed, for readability. **Red**: changes vs. the original VCF; highlight: errors.

##### 2.3.5 IUPACs

**From Table S9:**

| # Variants | Genozip | LiftoverVcf | CrossMap |
|---|---|---|---|
| 12 | ***Lifted*** | ***Variant Loss*** | ***Data Corruption*** |

**Genozip oSTATUS:** `OkRefSameSNPIupac`

The SNP test file has 12 variants at loci that contain a non-ACTGN IUPAC base in GRCh38. All 12 variants have a REF that is a base that is included in the mapped IUPAC base in GRCh38. For example (17, 81077361, T) is mapped to (chr17, 83129591, W). W is defined as A or T.

Since T is included in W, Genozip calls this variant as `OkRefSameSNP` and lifts it. LiftoverVcf rejects this variant because T≠W, which is a valid call but yet an unfortunate loss of data.

CrossMap on the other hand, replaces the REF with the IUPAC base, thereby generating variants that not only contain less information than the original variant (as the haplotypes with GT=0 had a definite base as specified by the original REF, not an ambiguous one) and hence represent a Data Loss, but are also noncompliant with the VCF 4.3 specification (violation of requirement 1.6.1-REF: "Each base must be one of A,C,G,T,N") and hence are likely to break downstream analysis tools:

```
#CHROM  POS        ID          REF  ALT
13      100973393  LN=3041971  K    T
13      100973395  LN=3041972  Y    T
17      83128871   LN=3550982  K    T
17      83128888   LN=3550984  Y    T
17      83129591   LN=3550985  W    A
17      83130798   LN=3550987  Y    T
17      83130998   LN=3550988  Y    T
17      83131245   LN=3550989  R    A
17      83131933   LN=3550990  Y    T
17      83133010   LN=3550991  R    A
17      83133390   LN=3550993  Y    T
17      83133686   LN=3550994  Y    T
```

#### 2.3.6 REF change, not to ALT, in bi-allelic SNPs when AF<1

**From Table S9:**

| # Variants | Genozip | LiftoverVcf | CrossMap |
|---|---|---|---|
| 68 | ***Variant Loss*** | ***Variant Loss*** | ***Data Corruption*** |

**Genozip oSTATUS:** `RefNewAlleleSNP`

When the REF base of a SNP changes between the references, and unless this is a REF⇄ALT switch in a bi-allelic SNP, CrossMap simply updates the new REF without updating ALT. This is correct only in the case where there are no haplotypes with the REF allele (i.e.

$\sum_{alt} AC_{alt} = AN$ or $\sum_{alt} AF_{alt} = 1$).

Example:

Original VCF (GRCh37):

```
#CHROM   POS        ID         REF      ALT     INFO(partial)
1        13808732   LN=20854   C        T       AC=1;AF=0.500;AN=2
```

CrossMap incorrectly-lifted VCF (GRCh38):

```
#CHROM   POS        ID         REF      ALT     INFO(partial)
1        13482278   LN=20854   G        T       AC=1;AF=0.500;AN=2
```

Since G is a new allele the correct lifting should have been:

```
#CHROM   POS        ID         REF      ALT     INFO(partial)
1        13482278   LN=20854   G        T,C     AC=1,1;AF=0.5,0.5;AN=2
```

Genozip and LiftoverVcf, in contrast, reject these variants as they cannot handle adding an allele.

##### 2.3.7 REF change in multi-allelic SNPs when AF<1

**From Table S9:**

| # Variants | Genozip | LiftoverVcf | CrossMap |
|---|---|---|---|
| 30 | ***Variant Loss*** | ***Variant Loss*** | ***Data Corruption*** |

**Genozip oSTATUS:** `RefMultiAltSwitchSNP`

In cases of multi-allelic SNPs with a reference base change, CrossMap changes REF without updating ALT, regardless of whether the new reference is one of the ALT alleles.

Example:

Original VCF (GRCh37):

```
#CHROM   POS        ID          REF     ALT   INFO(partial)
18       77831522   LN=3663131  G       C,T   AC=1,1;AF=0.500,0.500;AN=2
```

CrossMap incorrectly-lifted VCF (GRCh38):

```
#CHROM   POS        ID          REF     ALT   INFO(partial)
18       80073165   LN=3663131  C       C,T   AC=1,1;AF=0.500,0.500;AN=2
```

This correct lifting would have been a REF⇄ALT switch:

```
#CHROM   POS        ID          REF     ALT   INFO(partial)
18       80073165   LN=3663131  C       G,T   AC=0,1;AF=0,0.500;AN=2
```

Genozip and LiftoverVcf, in contrast, reject these variants as they cannot handle REF changes in multi-allelic SNPs.

#### 2.3.8 REF change, not to ALT, in bi-allelic SNPs when AF=1

**From Table S9:**

| # Variants | Genozip | LiftoverVcf | CrossMap |
|---|---|---|---|
| 47 | ***Lifted*** | ***Variant Loss*** | ***Lifted*** |

**Genozip oSTATUS:** `OkNewRefSNP`

In these bi-allelic SNP variants, there is a reference base change, and the sample doesn't contain any haplotypes with the REF allele, i.e. AC=AN or AF=1. Therefore, it is permissible to just update the REF (the old REF that would normally become one of the ALT alleles, is redundant in this case since it has AF=0). However, LiftoverVcf fails to do so, needlessly rejecting these variants.

Example:

Original VCF (GRCh37):

```
#CHROM   POS        ID          REF      ALT   INFO(partial)
1        99597759   LN=137361   C        G     AC=2;AF=1.00;AN=2
```

Genozip and CrossMap correctly-lifted VCF (GRCh38)—REF replacement OK if AF=1

```
#CHROM   POS        ID           REF      ALT   INFO(partial)
chr1     99132203   LN=137361    A        G     AC=2;AF=1.00;AN=2
```

##### 2.3.9 REF⇄ALT switch proportions

We now turn our attention to the 0.7% (29,635 out of 4,109,729) of the variants in our test file that are categorized as REF⇄ALT switches. These variants are dropped by CrossMap and in LiftoverVcf, they are either dropped or some of their annotations are dropped, depending on the command line options used. In contrast. Genozip lifts them over correctly, updating the annotations that sensitive to REF⇄ALT switches (see https://genozip.com/dvcf-rendering.html)

Here we show that despite the overall number of these variants being relatively small, they are not uniformly distributed across the genome but rather preferentially located in certain regions, and therefore dropping them may introduce bias in certain downstream analyses, such as GWAS or selection scans.

We divided the genome into 30,749 *windows* of 100 kb each, and counted for each 100-kb window the total number of variants in the SNP test file within the window vs. the number of those variants that are a REF⇄ALT switch. This was done by leveraging Genozip's internal genome-wide position called GPOS (for Global POSition). The position is that of the base as it appears in the original FASTA file used to generate the reference file, when all contigs are concatenated in the order they appear in the FASTA.
The first command, below, outputs the number of variants per 100-kb window in the test file. The first column in the output is the count of variants in a 100-kb window and the second is the sequential GPOS in units of 100k (the first line missing a number is window 0). For brevity, we show here the first 6 lines.

```
> genocat snp-37-38.d.vcf.genozip -e hs37d5.ref.genozip --gpos
-HG | cut -f2 | rev | cut -c6- | rev | uniq -c
    113
     60 1
     79 2
      4 3
      6 4
    124 5
```

The second command, below, outputs the number of variants with REF⇄ALT switch per 100-kb window in the test file. The first column of the output is the count of variants with REF⇄ALT switch and the second is the sequential GPOS in units of 100k. For brevity, we show here first 6 regions that have any REF⇄ALT switch variants at all):

```
> genocat snp-37-38.d.vcf.genozip -e hs37d5.ref.genozip --gpos
-HG --show-dvcf --grep OkRefAltSwitchSNP | cut -f4 | rev |cut
-c6- | rev | uniq -c
      1 7
      2 8
      1 11
      1 12
      1 13
     22 15
```

The proportion of REF⇄ALT switch variants compared to the total number of variants for each 100-kb window of our SNP test file are shown in Figure 1C. It is easy to see that distribution of REF⇄ALT switch variants across the genome is highly non-uniform – the vast majority of the 100kb windows have very few REF⇄ALT switch variants, while a small number of windows have a very high percentage of REF⇄ALT switch variants – some in which over 80% of the variants are switches. Therefore, when CrossMap or LiftoverVcf drop all variants with a REF⇄ALT switch, they are potentially introducing bias to the data that might impact downstream analyses. Another view of the same data is presented in Figure S3.

We then proceeded to compare the GRCh38 coordinates of REF⇄ALT switch variants to the regions of the GRCh38 reference genome with known issues, downloaded from https://ftp.ncbi.nlm.nih.gov/pub/grc/human/GRC/Issue_Mapping/GRCh38.p13_issues.gff3. Indeed, there is significant overlap between the loci with REF⇄ALT switches and regions of the reference genome known to be problematic, see Figure S4. The R script used for this analysis and generate the Figure S4 is available from https://github.com/divonlan/genozip-dvcf-results/tree/main/Fig-2A.

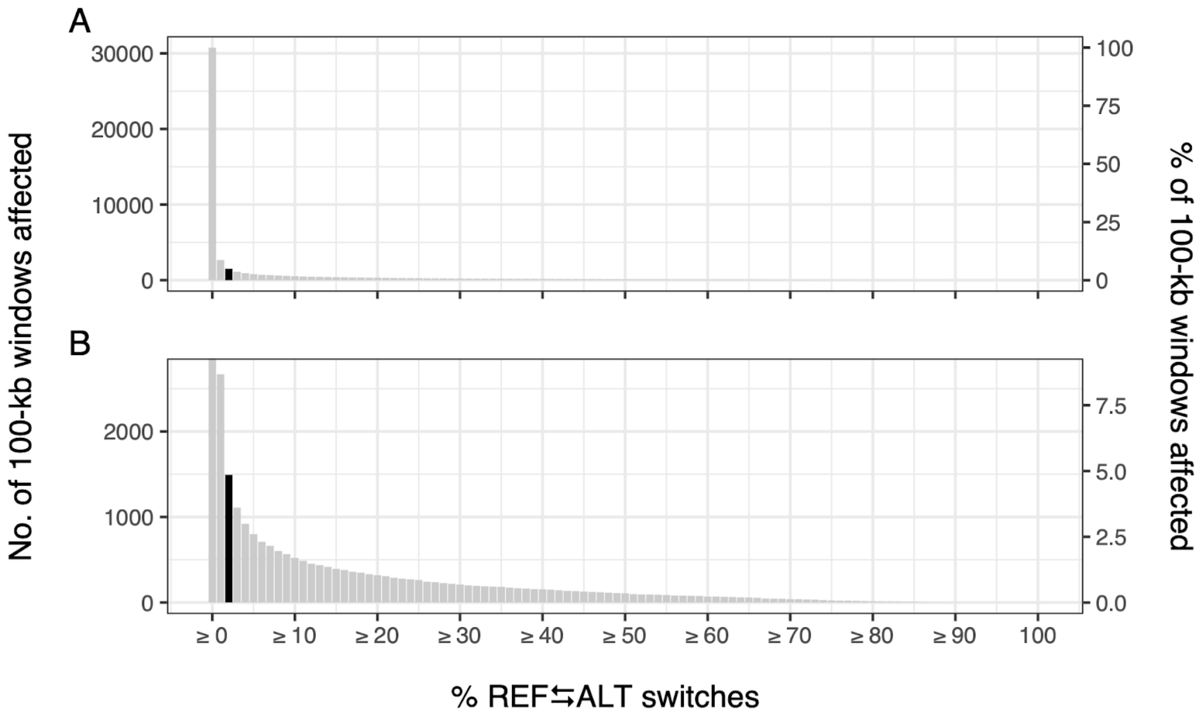

**Figure S3. Distribution of the number of 100-kb windows with at least *x*% REF⇄ALT switch variants**. A. The number of 100-kb windows was calculated for increments of 1% of REF⇄ALT switch variants content. The y-axis on the right side of the figure indicates the corresponding percentage of affected windows relative to all 100-kb windows. B. A close up of the distribution showing that nearly 5% (black bar) of all 100-kb windows in the human genome contain at least 2% REF⇄ALT switch variants amongst all variants in the window.

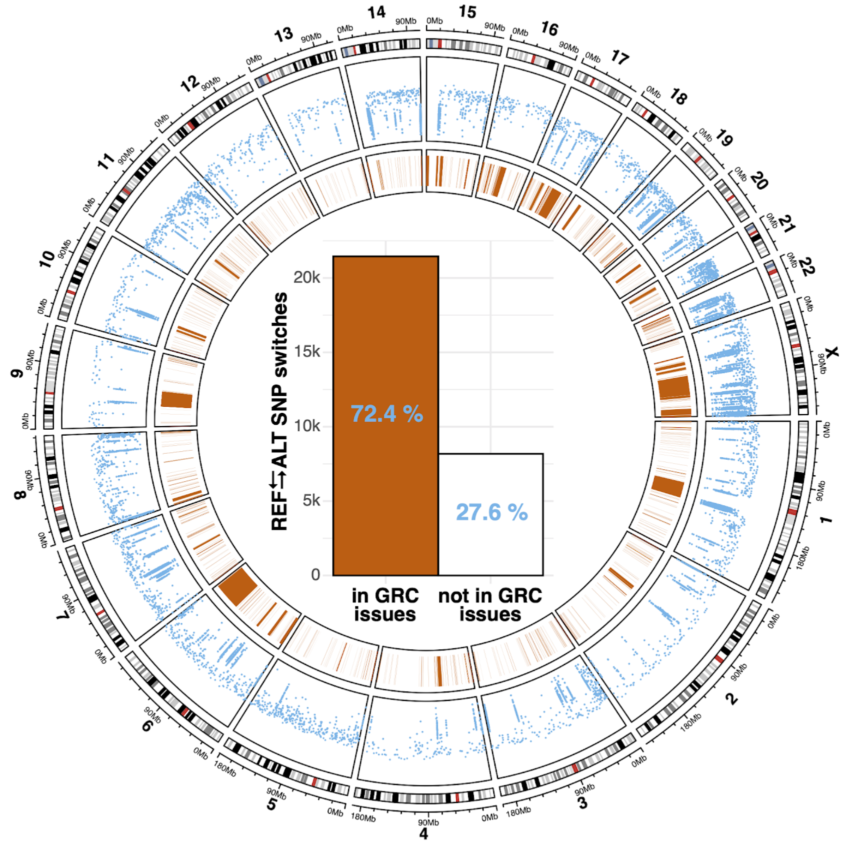

**Figure S4.** Distribution and functional impact of REF⇄ALT allele switches in SNP variants. Circos plot: the location of REF⇄ALT allele switches are shown in the blue rainfall plot, with GRC-identified problematic regions shown as orange polygons. Bar plot: Number (bars) and percentage (blue text) of REF⇄ALT allele switches inside or outside problematic GRC regions.

#### 2.4 Benchmark summary

**Table S12**: Summary of correctly (in green: Lifted or Unmapped) vs incorrectly (in red: Data Corruption or Variant Loss) lifted variants for each tested tool. For each lifting tool, we show the number of variants and the percentage of variants falling under each major category, including outcomes that could negatively impact downstream analyses (i.e., Data Corruption).

| | *Indels* | | | | | | *SNPs* | | | | | |
|---|---|---|---|---|---|---|---|---|---|---|---|---|
| | *Genozip* | | *LiftoverVcf* | | *CrossMap* | | *Genozip* | | *LiftoverVcf* | | *CrossMap* | |
| *Total* | 18706 | 100% | 18706 | 100% | 18706 | 100% | 4109729 | 100% | 4109729 | 100% | 4109729 | 100% |
| *Correct* | 18663 | 99.8% | 18304 | 97.9% | 18279 | 97.7% | 4109631 | 99.998% | 4064248 | 98.9% | 4064295 | 98.9% |
| *Correct that are lifted over* | 18565 | 99.2% | 18206 | 97.3% | 18201 | 97.3% | 4082903 | 99.3% | 4037520 | 98.2% | 4037567 | 98.2% |
| *Incorrect* | 43 | 0.2% | 402 | 2.1% | 427 | 2.3% | 98 | 0.002% | 45481 | 1.1% | 45434 | 1.1% |
| *Incorrect that lead to Data Corruption* | 0 | 0% | 306 | 1.6% | 356 | 1.9% | 0 | 0% | 15689 | 0.4% | 15799 | 0.4% |
| *Incorrect that are REF⇄ALT switches* | 0 | 0% | 257 | 1.3% | 257 | 1.3% | 0 | 0% | 29635 | 0.7% | 29635 | 0.7% |

### Section 3: ClinVar analysis

#### 3.1 Data preparation

We analysed the ClinVar file from the week of 03 Jan 2022 downloaded from https://ftp.ncbi.nlm.nih.gov/pub/clinvar/vcf_GRCh37/weekly/clinvar_20220103.vcf.gz

The analysis can be run by executing the bash script `run-clinvar-37-38.sh` in the github repository https://github.com/divonlan/genozip-dvcf-results. The key parts of this script are:

Lifting the file to a DVCF:

```
> genozip --echo --chain
shared/GRCh37_to_GRCh38.matched.chain.genozip --add-line-numbers
--match-chrom-to-reference shared/clinvar.37.vcf.gz -o
clinvar-37-38/clinvar-37-38.d.vcf.genozip
```

As before, to allow easy detection of variants with potential issues, we created a ClinVar GRCh37 single-coordinate file that contains an extra `INFO/oSTATUS` field, as well as the line numbers in the ID field:

```
> genocat clinvar-37-38/clinvar-37-38.d.vcf.genozip --single -o
clinvar-37-38/clinvar.37.annotated.vcf --show-ostatus
```

For CrossMap, we used the following command:

```
> CrossMap.py vcf shared/GRCh37_to_GRCh38.matched.chain
clinvar-37-38/clinvar.37.annotated.vcf
shared/GRCh38_full_analysis_set_plus_decoy_hla.fa.gz
clinvar-37-38/clinvar.38.CrossMap.vcf
```

For LiftoverVcf, we used the following command:

```
> gatk --java-options '-Xmx16g -XX:ParallelGCThreads=1' LiftoverVcf
--INPUT clinvar-37-38/clinvar.37.annotated.vcf --OUTPUT
clinvar-37-38/clinvar.38.gatk.vcf --CHAIN
shared/GRCh37_to_GRCh38.matched.chain --REJECT
clinvar-37-38/clinvar.38.gatk.rejects.vcf --RECOVER_SWAPPED_REF_ALT
--REFERENCE_SEQUENCE
shared/GRCh38_full_analysis_set_plus_decoy_hla.fa.gz
--TAGS_TO_REVERSE AF_ESP --TAGS_TO_REVERSE AF_EXAC --TAGS_TO_REVERSE
AF_TGP --TAGS_TO_DROP DUMMY
```

#### 3.2 Genozip analysis

Genozip has no variants with corrupted data, however it did drop 201 variants (133 SNPs, 68 non-SNPs) which have a valid mapping in the chain file, because the allele represented by the GRCh38 reference is neither the REF nor the ALT allele (it also *correctly* dropped 1428 variants because RefNotMappedInChain and RefSplitInChain).

**Table S13:** ClinVar variants dropped by Genozip despite having a mapping in the chain file. This occurs because the allele in the GRCh38 (Luft) reference is a new allele, neither REF nor ALT. This does not include variants correctly dropped due to lack of mapping in the chain file.

| Type | # | Issue | ALLELEID | CLNSIG |
|---|---|---|---|---|
| Data loss | 201 | Allele in GRCh38 is a new allele (neither REF nor ALT) | 696043,799115,685731,15474,964796,1170574,696055,696058,1167865,581994,696139,15681,696267,696378,696404,447571,425007,392854,697512,286118,451010,286231,390508,221328,708391,1154227,980680,963007,173441,697258,1154060,698145,452738,453319,215271,33220,455066,698872,174523,54118,1037727,1037728,226789,700218,389801,710823,961851,699897,699899,699900,699901,699902,699903,711517,790800,1155979,459012,959873,700419,700425,700426,700428,700429,700431,700436,700482,1008398,1156206,851221,174261,174548,700865,700867,1162096,800875,1162094,1162095,487407,549956,700869,700871,700872,692808,701351,397737,622275,701488,47952,240741,53156,701260,20181,167056,712775,390514,33351,838853,1156807,1166021,167084,917805,244624,150504,1156610,712655,702499,175552,175851,702467,713776,461651,702807,702808,703029,167190,529374,703257,703335,703373,242133,1157890,465467,818455,468332,1033832,343827,789384,1157991,376201,375315,706355,704364,704365,715791,55368,705368,245748,245880,245942,964061,536982,176732,705090,431872,334260,797925,682788,963904,1169844,882486,882487,40517,1158756,684879,684881,1158835,705475,705477,680034,966169,705738,963022,390511,390657,390590,390591,963023,705823,439050,38485,31934,47980,706250,706252,26660,706257,706259,706260,706263,706277,243824,963026,706035,670963,792118,706116,549821,670737,706133,706162,706163,706164,706168,706169,706170,706171,982298,472107,706177,706179,706180 | 135 Benign<br>17 Uncertain significance<br>15 Likely benign<br>10 Pathogenic<br>6 Likely pathogenic<br>6 Benign/Likely benign<br>3 not provided<br>3 association<br>2 risk factor<br>2 Conflicting interpretations of pathogenicity<br>1 other<br>1 drug response |

### 3.3 CrossMap analysis

Of the 969,410 variants in the file, 967,781 were lifted, and 1628 failed to lift. Some of the lifted variants were corrupt, and some of the dropped variants were unnecessarily dropped.

**Table S14:** ClinVar variants incorrectly dropped by CrossMap (excluding variants correctly dropped due to lack of mapping in the chain file), and variants lifted incorrectly.

| Type | # | Issue | ALLELEID | CLNSIG |
|---|---|---|---|---|
| Data loss | 204 | Failed to lift REF⇄ALT switch | 177885,191721,389423,106000,1163377,1163378,862175,1153237,353057,227743,1164070,1153536,249770,655075,1153962,389508,177658,141535,102135,389649,390440,670179,671101,291712,291961,789736,193757,389612,251402,1168025,167900,251996,36716,177851,54119,1037729,141976,141771,178204,141770,390476,390568,227764,389793,18435,699898,683010,1155956,684027,662663,662667,136076,1171823,304611,390475,313496,1156147,1156207,174544,54805,700866,800873,1171929,143080,654535,1171932,790945,701352,701350,701349,701348,701347,270003,142604,53180,389879,389938,254076,175704,254276,389897,17210,137366,701599,140343,54569,54553,254491,54551,55703,1157066,190700,254805,791396,1157289,230590,1157471,339619,323041,873957,656305,433550,390078,132224,192275,1157931,1158130,256553,344888,1087130,345325,1173004,791785,269599,506140,1163638,1173059,329799,256613,390307,791872,508906,344604,230979,257300,1168415,433895,142630,231042,257199,334255,177983,1173294,169607,344231,349365,178100,45542,353525,137338,257355,257356,257358,23429,716929,1158836,351613,1169888,106610,1164570,1164571,1164572,1164573,792021,817902,508193,817919,817923,817930,508196,818000,818032,818037,818051,818053,818054,818055,818056,818057,818058,818060,818090,818094,818115,818122,717806,1159714,1159720,1159723,101448,792471,352903,134697,99042,98295,52521,800324,52519,656729,24924,670821,669859,25543,99678,101226,45455,352797,1159494,671190,352799,99399,25405,339086,99598 | 160 Benign<br>21 drug response<br>9 Likely benign<br>8 Benign/Likely benign<br>3 Conflicting interpretations of pathogenicity<br>2 Uncertain significance<br>1 Pathogenic |

**Table S14:** (continued from previous page)

| Type | # | Issue | ALLELEID | CLNSIG |
|---|---|---|---|---|
| Data Corruption | 201 | Lifted has bad REF field | 696043,799115,685731,15474,964796,1170574,696055,696058,1167865,581994,696139,15681,696267,696378,696404,447571,425007,392854,697512,286118,451010,286231,390508,221328,708391,1154227,980680,963007,173441,697258,1154060,698145,452738,453319,215271,33220,455066,698872,174523,54118,1037727,1037728,226789,700218,389801,710823,961851,699897,699899,699900,699901,699902,699903,711517,790800,1155979,459012,959873,700419,700425,700426,700428,700429,700431,700436,700482,1008398,1156206,851221,174261,174548,700865,700867,1162096,800875,1162094,1162095,487407,549956,700869,700871,700872,692808,701351,397737,622275,701488,47952,240741,53156,701260,20181,167056,712775,390514,33351,838853,1156807,1166021,167084,917805,244624,150504,1156610,712655,702499,175552,175851,702467,713776,461651,702807,702808,703029,167190,529374,703257,703335,703373,242133,1157890,465467,818455,468332,1033832,343827,789384,1157991,376201,375315,706355,704364,704365,715791,55368,705368,245748,245880,245942,964061,536982,176732,705090,431872,334260,797925,682788,963904,1169844,882486,882487,40517,1158756,684879,684881,1158835,705475,705477,680034,966169,705738,963022,390511,390657,390590,390591,963023,705823,439050,38485,31934,47980,706250,706252,26660,706257,706259,706260,706263,706277,243824,963026,706035,670963,792118,706116,549821,670737,706133,706162,706163,706164,706168,706169,706170,706171,982298,472107,706177,706179,706180 | 135 Benign<br>17 Uncertain significance<br>15 Likely benign<br>10 Pathogenic<br>6 Likely pathogenic<br>6 Benign/Likely benign<br>3 not provided<br>3 association<br>2 risk factor<br>1 other<br>1 drug response<br>2 Conflicting interpretations of pathogenicity |
| Data Corruption | 4 | Lifted failed to switch REF and ALT | 776998,657462,198367,438998 | 3 Benign<br>1 Likely benign |
| Data Corruption | 4 | Mapped despite no mapping in chain file | 1097855,407586,1156794,1157524 | 2 Likely benign<br>2 Benign |

#### 3.4 LiftoverVcf analysis

Of the 969,410 variants in the file, 967,816 were lifted, and 1594 failed to lift. Some of the lifted variants were corrupt, and some of the dropped variants were unnecessarily dropped:

**Table S15:** ClinVar variants incorrectly dropped by LiftoverVcf (excluding variants correctly dropped due to lack of mapping in the chain file), and variants lifted incorrectly.

| Type | # | Issue | ALLELEID | CLNSIG |
|---|---|---|---|---|
| Data loss | 162 | Allele in GRCh38 is a new allele (neither REF nor ALT) | 696043,685731,15474,964796,1170574,696055,696058,581994,696139,15681,696267,696378,696404,447571,425007,392854,697512,451010,708391,173441,697258,698145,452738,453319,33220,455066,698872,174523,54118,1037728,226789,700218,389801,710823,699897,699899,699900,699901,699902,699903,711517,459012,959873,700419,700425,700426,700428,700429,700431,700436,700482,1008398,1156206,851221,174261,174548,700865,700867,1162096,800875,1162094,1162095,700869,700871,700872,692808,701351,397737,622275,701488,47952,240741,53156,701260,20181,167056,712775,33351,838853,1166021,167084,917805,150504,712655,702499,175552,175851,702467,713776,702807,702808,167190,529374,703257,703335,703373,242133,465467,818455,468332,1033832,343827,789384,376201,375315,706355,704364,704365,715791,705368,245748,245880,245942,536982,705090,431872,334260,797925,682788,963904,882486,882487,40517,684879,684881,705475,705477,680034,966169,705738,963022,390591,705823,38485,31934,47980,706250,706252,26660,706257,706259,706260,706263,706277,243824,963026,706035,706116,549821,706133,706162,706163,706164,706168,706169,706170,706171,982298,472107,706177,706179,706180 | 107 Benign<br>13 Uncertain significance<br>11 Likely benign<br>9 Pathogenic<br>5 Likely pathogenic<br>5 Benign/Likely benign<br>3 not provided<br>3 association<br>2 risk factor<br>2 Conflicting interpretations of pathogenicity<br>1 other<br>1 drug response |
| Data loss | 4 | Failed to lift REF⇄ALT switch of non-SNPs | 1037729,178204,683010,800324,16 | 2 Uncertain significance<br>2 Benign |
| Data corruption | 39 | Incorrect REF field (GRCh38 is a new allele) | 964796,581994,1037728,389801,959873,851221,167056,167084,167190,818455,789384,245748,245880,245942,536982,431872,797925,963904,966169,963022,26660,963026 | 28 Benign<br>4 Uncertain significance<br>4 Likely benign<br>1 Pathogenic<br>1 Likely pathogenic<br>1 Benign/Likely benign |
| Data corruption | 4 | Lifted failed to switch REF and ALT | 1037729,683010 | 3 Benign<br>1 Likely benign |

#### 3.5 ClinVar benchmark – summary

All tools dropped variants of “Pathogenic” clinical significance, leading to the conclusion that lift over techniques should never be used in a clinical setting. However, CrossMap and LiftoverVcf did worse than that – they also *incorrectly* lifted 10 (CrossMap) or 1 (LiftoverVcf) variants of “Pathogenic” clinical significance, resulting in a corrupt REF field of these variants, that could potentially lead to incorrect clinical diagnosis.

### Section 4: GRCh38 and Telomere-to-Telomere

We repeated the tests described in section 1 and 3, but this time between the GRCh38 and Telomere-to-Telomere v1.0 reference.

- For the SNP test file, we used the GRCh38 file which was the output of the Genozip lift-over from GRCh37 to GRCh38.

- For the indels test file, we extracted the indel variants from a GRCh38 version of the 1000 Genome Project. This is an independent analysis of the 1KGP data, and *not* a lift-over of the GRCh37 file we used for testing. The GRCh38 file contains significantly more indel variants. The file was obtained from: ftp://ftp.1000genomes.ebi.ac.uk/vol1/ftp/data_collections/1000G_2504_high_coverage/working/20201028_3202_raw_GT_with_annot/

- For the ClinVar test file, we obtained the GRCh38 version of the same data used in section 3, from: ftp://ftp.ncbi.nlm.nih.gov/pub/clinvar/vcf_GRCh38/weekly/clinvar_20210724.vcf.gz

- The GRCh38 to T2Tv1.0 chain file was obtained from: http://t2t.gi.ucsc.edu/chm13/hub/t2t-chm13-v1.0/hg38Lastz/hg38.t2t-chm13-v1.0.over.chain.gz

- The T2Tv1.0 reference file was obtained from: https://s3-us-west-2.amazonaws.com/human-pangenomics/T2T/CHM13/assemblies/chm13.draft_v1.0.fasta.gz

- The GRCh38 reference, was obtained from: ftp://ftp.1000genomes.ebi.ac.uk/vol1/ftp/technical/reference/GRCh38_reference_genome/GRCh38_full_analysis_set_plus_decoy_hla.fa

As before, we provide bash scripts which execute the entire analysis including downloading the files. These scripts also serve as precise instructions for reproducing our results. They can be found in https://github.com/divonlan/genozip-dvcf-results.

We observe that the mapping of GRCh38 to T2T is significantly more complex than the mapping of GRCh37 to GRCh38, by the measure of the number of unique alignments in their respective chain files:

```
> genocat --show-chain GRCh37_to_GRCh38.matched.chain.genozip | wc -l
18389

> genocat --show-chain hg38.t2t-chm13-v1.0.over.matched.chain.genozip|wc -l
865173
```

A mapping of how each of the three tools handles the various variant categories appears in Tables S18, S19 and S20. We note that the tools diverge significantly with regards to how they handle the various cases. Note that “lifted” merely means that the variant appears in the output file—not that it is correct. In fact, there are many known cases in which the variant is incorrect—these are documented in Sections 2.2 and 2.3 and a mapping to the relevant section appears in the second column of each table. We also noted some new categories that did not appear in the GRCh37-to-GRCh38 case; these are marked as “New”. We did not perform a detailed analysis of the causes for the differences between the tools in this case of GRCh38-to-T2T liftover, which should be the purpose of future research.

**Table S18**. Categorization of indel variants and how each tool handles problematic variants.

| Genozip oStatus | Section | Count | Genozip | LiftoverVcf | CrossMap |
|---|---|---|---|---|---|
| **OkRefSameIndel** | N/A | 155,695 | Lifted | Lifted | Lifted |
| **OkRefSameNotLeftAnc** | N/A | 30,654 | Lifted | 30123 Lifted<br>531 Dropped | Lifted |
| **OkRefSameDelRev** | 2.2.8 | 3,823 | Lifted | 3535 Lifted<br>288 Dropped | Lifted |
| **OkRefSameInsRev** | 2.2.7 | 2,010 | Lifted | 1998 Lifted<br>12 Dropped | Lifted |
| **OkRefAltSwitchIndelRpts** | 2.2.3 | 1,738 | Lifted | 1735 Lifted<br>3 Dropped | Lifted |
| **OkRefSameNDNIRev** | New | 1,575 | Lifted | 1271 Lifted<br>304 Dropped | Lifted |
| **OkRefAltSwitchWithGap** | 2.2.5 | 871 | Lifted | Variant dropped | Variant dropped |
| **OkRefAltSwitchIndelFlank** | 2.2.3 | 389 | Lifted | Lifted | Lifted |
| **RefSplitInChain** | 2.2.6 | 12,147 | Dropped | Dropped | Lifted 2554<br>Dropped 9593 |
| **RefNotMappedInChain** | N/A | 5,688 | Dropped | Dropped | Lifted 2127<br>Dropped 3561 |
| **RefMultiAltSwitchIndel** | New | 3,002 | Dropped | Lifted 1695<br>Dropped 1307 | Lifted |
| **RefNewAlleleNotLeftAnc** | New | 2,835 | Dropped | Lifted 53<br>Dropped 2782 | Lifted |
| **RefNewAlleleNDNI** | New | 2,459 | Dropped | Lifted 44<br>Dropped 2415 | Lifted |
| **RefNewAlleleDelRefChanged** | 2.2.9 | 1,429 | Dropped | Lifted 66<br>Dropped 1363 | Lifted |
| **RefNewAlleleInsSameRef** | 2.2.10 | 1,181 | Dropped | Lifted | Lifted |
| **RefNewAlleleDelSameRef** | 2.2.10 | 512 | Dropped | Lifted 475<br>Dropped 37 | Lifted |
| **RefNewAlleleIndelNoSwitch** | 2.2.11 | 380 | Dropped | Lifted | Lifted |
| **RefNewAllelInsRefChanged** | 2.2.9 | 313 | Dropped | Lifted 121<br>Dropped 192 | Lifted |
| **INFO/AF** | New | 167 | Dropped | Dropped | Lifted |
| **Total** | | **226,868** | | | |

**Table S19**. Categorization of SNP variants and how each tool handles problematic variants.

| Genozip oStatus | Section | Count | Genozip | LiftoverVcf | CrossMap |
|---|---|---|---|---|---|
| **OkRefSameSNP** | N/A | 2,288,044 | Lifted | Lifted | Lifted |
| **OkRefAltSwitchSNP** | 2.3.3 | 1,705,586 | Lifted | INFO/AC dropped | Dropped |
| **OkRefSameSNPRev** | 2.3.4 | 22,266 | Lifted | Lifted | Lifted |
| **OkNewRefSNP** | 2.3.8 | 1,869 | Lifted | Dropped | Lifted |
| **NoMappingInChainFile** | N/A | 61,771 | Dropped | Dropped | Dropped |
| **RefNewAlleleSNP** | 2.3.6 | 1,934 | Dropped | Dropped | Lifted |
| **RefMultiAltSwitchSNP** | 2.3.7 | 1,433 | Dropped | Dropped | Lifted |
| **Total** | | **4,060,637** | | | |

**Table S20**. Categorization of ClinVar variants and how each tool handles problematic variants.

| Genozip oStatus | Section | Count | Genozip | LiftoverVcf | CrossMap |
|---|---|---|---|---|---|
| **OkRefSameSNP** | N/A | 857,612 | Lifted | Lifted | Lifted |
| **OkRefSameIndel** | N/A | 86,095 | Lifted | Lifted | Lifted |
| **OkRefAltSwitchSNP** | 2.3.3 | 11,055 | Lifted | Lifted | Dropped |
| **OkRefSameNotLeftAnc** | N/A | 5,972 | Lifted | Lifted | Lifted |
| **OkRefSameSNPRev** | 2.3.4 | 2,058 | Lifted | Lifted | Lifted |
| **OkRefAltSwitchIndelRpts** | 2.2.3 | 734 | Lifted | Lifted | Lifted |
| **OkRefAltSwitchWithGap** | 2.2.5 | 432 | Lifted | Dropped | Dropped 431<br>Lifted 1 |
| **OkRefAltSwitchIndelFlank** | 2.2.3 | 152 | Lifted | Lifted | Lifted |
| **OkRefSameDelRev** | 2.2.8 | 89 | Lifted | Lifted 87<br>Dropped 2 | Lifted |
| **OkRefAltSwitchDelToIns** | 2.2.4 | 53 | Lifted | Dropped | Lifted |
| **OkRefAltSwitchNotLeftAnc** | New | 50 | Lifted | Dropped | Dropped 34<br>Lifted 16 |
| **OkRefSameInsRev** | 2.2.7 | 48 | Lifted | Lifted | Lifted |
| **RefNewAlleleInsSameRef** | 2.2.10 | 1,071 | Dropped | Lifted | Lifted |
| **RefSplitInChain** | 2.2.6 | 1,003 | Dropped | Dropped | Dropped 897<br>Lifted 106 |
| **RefNotMappedInChain** | N/A | 859 | Dropped | Dropped | Dropped 791<br>Lifted 68 |
| **RefNewAlleleSNP** | 2.3.6 | 732 | Dropped | Dropped | Lifted |
| **RefNewAlleleDelRefChanged** | 2.2.9 | 520 | Dropped | Dropped | Lifted |
| **RefNewAlleleNotLeftAnc** | New | 457 | Dropped | Dropped | Lifted 457<br>Dropped 1 |
| **RefNewAlleleDelSameRef** | 2.2.10 | 340 | Dropped | Lifted | Lifted |
| **RefNewAllelInsRefChanged** | 2.2.9 | 111 | Dropped | Dropped 109<br>Lifted 2 | Lifted |
| **RefNewAlleleIndelNoSwitch** | 2.2.11 | 42 | Dropped | Lifted | Lifted |
| **ChromNotInPrimReference** | N/A | 1 | Dropped | Dropped | Dropped |
| **Total** | | **969,486** | | | |

### Appendix: Analysis script output

Our analysis scripts in the https://github.com/divonlan/genozip-dvcf-results repository generate analysis output files for each tool. The Genozip analysis file summarizes the number of variants assigned to each category, while the analysis files for CrossMap and LiftoverVcf summarize the number of variants in each Genozip category, that were either lifted or dropped by the tool.

We therefore have analysis files for each of the three tools (Genozip, CrossMap and LiftoverVcf), for each of the three tests (Indels, SNPs and ClinVar) for each of the two datasets (GRCh37->GRCh38 and GRCh38->T2T):

1) Indels – GRCh37 to GRCh38

```
Genozip
Showing counts of o$TATUS (did_i=17). Total items=18706 Number of
categories=14
OkRefSameIndel  18201 97.30%
OkRefAltSwitchIndelRpts     153   0.82%
RefNotMappedInChain   78    0.42%
OkRefAltSwitchWithGap 71    0.38%
OkRefSameDelRev  67    0.36%
OkRefSameInsRev  40    0.21%
OkRefAltSwitchIndelFlank    27    0.14%
RefSplitInChain  20    0.11%
RefNewAlleleDelRefChanged  13    0.07%
RefNewAlleleInsSameRef      11    0.06%
RefNewAlleleIndelNoSwitch  9     0.05%
OkRefAltSwitchDelToIns      6     0.03%
RefNewAlleleInsRefChanged  6     0.03%
RefNewAlleleDelSameRef      4     0.02%

CrossMap
data=indel primary=37 luft=38 tool=CrossMap
Lifted OkRefSameIndel: 18201
Lifted OkRefAltSwitchIndelRpts: 153
Lifted RefNotMappedInChain: 2
Failed RefNotMappedInChain: 76
Failed OkRefAltSwitchWithGap: 71
Lifted OkRefSameDelRev: 67
Lifted OkRefSameInsRev: 40
Lifted OkRefAltSwitchIndelFlank: 27
Lifted RefSplitInChain: 18
Failed RefSplitInChain: 2
Lifted RefNewAlleleDelRefChanged: 13
Lifted RefNewAlleleInsSameRef: 11
Lifted RefNewAlleleIndelNoSwitch: 9
Lifted OkRefAltSwitchDelToIns: 6
Lifted RefNewAlleleInsRefChanged: 6
Lifted RefNewAlleleDelSameRef: 4

LiftoverVcf
data=indel primary=37 luft=38 tool=gatk
Lifted OkRefSameIndel: 18201
Lifted OkRefAltSwitchIndelRpts: 153
Failed RefNotMappedInChain: 78
Failed OkRefAltSwitchWithGap: 71
Lifted OkRefSameDelRev: 67
Lifted OkRefSameInsRev: 40
Lifted OkRefAltSwitchIndelFlank: 27
Failed RefSplitInChain: 20
Failed RefNewAlleleDelRefChanged: 13
Lifted RefNewAlleleInsSameRef: 11
Lifted RefNewAlleleIndelNoSwitch: 9
Failed OkRefAltSwitchDelToIns: 6
Failed RefNewAlleleInsRefChanged: 6
Lifted RefNewAlleleDelSameRef: 4
```

2) SNPs – GRCh37 to GRCh38

```
Genozip
Showing counts of o$TATUS (did_i=17). Total items=4109729 Number of
categories=8
OkRefSameSNP     4037520    98.24%
OkRefAltSwitchSNP     29635 0.72%
RefNotMappedInChain   26728 0.65%
OkRefSameSNPRev 15689 0.38%
RefNewAlleleSNP 68    0.00%
OkNewRefSNP     47    0.00%
RefMultiAltSwitchSNP  30    0.00%
OkRefSameSNPIupac     12    0.00%

CrossMap
data=snp primary=37 luft=38 tool=CrossMap
Lifted OkRefSameSNP: 4037520
Failed OkRefAltSwitchSNP: 29635
Failed RefNotMappedInChain: 26728
Lifted OkRefSameSNPRev: 15689
Lifted RefNewAlleleSNP: 68
Lifted OkNewRefSNP: 47
Lifted RefMultiAltSwitchSNP: 30
Lifted OkRefSameSNPIupac: 12

LiftoverVcf
data=snp primary=37 luft=38 tool=gatk
Lifted OkRefSameSNP: 4037520
Lifted OkRefAltSwitchSNP: 29635
Failed RefNotMappedInChain: 26728
Lifted OkRefSameSNPRev: 15689
Failed RefNewAlleleSNP: 68
Failed OkNewRefSNP: 47
Failed RefMultiAltSwitchSNP: 30
Failed OkRefSameSNPIupac: 12
```

3) ClinVar – GRCh37 to GRCh38

```
Genozip
Showing counts of o$TATUS (did_i=17). Total items=969410 Number of
categories=19
OkRefSameSNP    870016    89.75%
OkRefSameIndel  90450 9.33%
OkRefSameNotLeftAnc   6557  0.68%
RefNotMappedInChain   1424  0.15%
OkRefSameSNPRev 501   0.05%
OkRefAltSwitchSNP     200   0.02%
RefNewAlleleSNP 133   0.01%
RefNewAlleleInsSameRef      36    0.00%
OkRefSameDelRev 33    0.00%
RefNewAlleleDelRefChanged  20    0.00%
OkRefSameInsRev 16    0.00%
RefNewAlleleNotLeftAnc      7     0.00%
OkRefAltSwitchIndelRpts     4     0.00%
RefSplitInChain 4     0.00%
OkRefAltSwitchWithGap 2     0.00%
OkRefAltSwitchNotLeftAnc    2     0.00%
RefNewAlleleInsRefChanged  2     0.00%
RefNewAlleleIndelNoSwitch  2     0.00%
RefNewAlleleDelSameRef      1     0.00%
```

```
CrossMap
data=clinvar primary=37 luft=38 tool=CrossMap
Lifted OkRefSameSNP: 870016
Lifted OkRefSameIndel: 90450
Lifted OkRefSameNotLeftAnc: 6557
Lifted RefNotMappedInChain: 2
Failed RefNotMappedInChain: 1422
Lifted OkRefSameSNPRev: 501
Failed OkRefAltSwitchSNP: 200
Lifted RefNewAlleleSNP: 133
Lifted RefNewAlleleInsSameRef: 36
Lifted OkRefSameDelRev: 33
Lifted RefNewAlleleDelRefChanged: 20
Lifted OkRefSameInsRev: 16
Lifted RefNewAlleleNotLeftAnc: 7
Lifted OkRefAltSwitchIndelRpts: 4
Lifted RefSplitInChain: 2
Failed RefSplitInChain: 2
Failed OkRefAltSwitchWithGap: 2
Failed OkRefAltSwitchNotLeftAnc: 2
Lifted RefNewAlleleInsRefChanged: 2
Lifted RefNewAlleleIndelNoSwitch: 2
Lifted RefNewAlleleDelSameRef: 1
```

```
LiftoverVcf
data=clinvar primary=37 luft=38 tool=gatk
Lifted OkRefSameSNP: 870016
Lifted OkRefSameIndel: 90450
Lifted OkRefSameNotLeftAnc: 6557
Failed RefNotMappedInChain: 1424
Lifted OkRefSameSNPRev: 501
Lifted OkRefAltSwitchSNP: 200
Failed RefNewAlleleSNP: 133
Lifted RefNewAlleleInsSameRef: 36
Lifted OkRefSameDelRev: 33
Failed RefNewAlleleDelRefChanged: 20
Lifted OkRefSameInsRev: 16
Failed RefNewAlleleNotLeftAnc: 7
Lifted OkRefAltSwitchIndelRpts: 4
Failed RefSplitInChain: 4
Failed OkRefAltSwitchWithGap: 2
Failed OkRefAltSwitchNotLeftAnc: 2
Failed RefNewAlleleInsRefChanged: 2
Lifted RefNewAlleleIndelNoSwitch: 2
Lifted RefNewAlleleDelSameRef: 1
```

4) Indels – GRCh38 to T2T

```
Genozip
Showing counts of o$TATUS (did_i=17). Total items=227096 Number of
categories=20
OkRefSameIndel	155695	68.56%
OkRefSameNotLeftAnc	30654	13.50%
RefSplitInChain	12147	5.35%
RefNotMappedInChain	5688	2.50%
OkRefSameDelRev	3823	1.68%
RefMultiAltSwitchIndel	3002	1.32%
RefNewAlleleNotLeftAnc	2835	1.25%
RefNewAlleleNDNI	2459	1.08%
OkRefSameInsRev	2010	0.89%
OkRefAltSwitchIndelRpts	1738	0.77%
OkRefSameNDNIRev	1575	0.69%
RefNewAlleleDelRefChanged	1429	0.63%
RefNewAlleleInsSameRef	1181	0.52%
OkRefAltSwitchWithGap	871	0.38%
RefNewAlleleDelSameRef	512	0.23%
OkRefAltSwitchIndelFlank	389	0.17%
RefNewAlleleIndelNoSwitch	380	0.17%
RefNewAlleleInsRefChanged	313	0.14%
OkRefAltSwitchDelToIns	228	0.10%
INFO/AF	167	0.07%

CrossMap
data=indel primary=38 luft=t2t tool=CrossMap
Lifted OkRefSameIndel: 155695
Lifted OkRefSameNotLeftAnc: 30654
Lifted RefSplitInChain: 2554
Failed RefSplitInChain: 9593
Lifted RefNotMappedInChain: 2127
Failed RefNotMappedInChain: 3561
Lifted OkRefSameDelRev: 3823
Lifted RefMultiAltSwitchIndel: 3002
Lifted RefNewAlleleNotLeftAnc: 2835
Lifted RefNewAlleleNDNI: 2459
Lifted OkRefSameInsRev: 2010
Lifted OkRefAltSwitchIndelRpts: 1738
Lifted OkRefSameNDNIRev: 1575
Lifted RefNewAlleleDelRefChanged: 1429
Lifted RefNewAlleleInsSameRef: 1181
Failed OkRefAltSwitchWithGap: 871
Lifted RefNewAlleleDelSameRef: 512
Lifted OkRefAltSwitchIndelFlank: 389
Lifted RefNewAlleleIndelNoSwitch: 380
Lifted RefNewAlleleInsRefChanged: 313
```

```
Lifted OkRefAltSwitchDelToIns: 228
Lifted INFO/AF: 167

LiftoverVcf
data=indel primary=38 luft=t2t tool=gatk
Lifted OkRefSameIndel: 155695
Lifted OkRefSameNotLeftAnc: 30123
Failed OkRefSameNotLeftAnc: 531
Failed RefSplitInChain: 12147
Failed RefNotMappedInChain: 5688
Lifted OkRefSameDelRev: 3535
Failed OkRefSameDelRev: 288
Lifted RefMultiAltSwitchIndel: 1695
Failed RefMultiAltSwitchIndel: 1307
Lifted RefNewAlleleNotLeftAnc: 53
Failed RefNewAlleleNotLeftAnc: 2782
Lifted RefNewAlleleNDNI: 44
Failed RefNewAlleleNDNI: 2415
Lifted OkRefSameInsRev: 1998
Failed OkRefSameInsRev: 12
Lifted OkRefAltSwitchIndelRpts: 1735
Failed OkRefAltSwitchIndelRpts: 3
Lifted OkRefSameNDNIRev: 1271
Failed OkRefSameNDNIRev: 304
Lifted RefNewAlleleDelRefChanged: 66
Failed RefNewAlleleDelRefChanged: 1363
Lifted RefNewAlleleInsSameRef: 1181
Failed OkRefAltSwitchWithGap: 871
Lifted RefNewAlleleDelSameRef: 475
Failed RefNewAlleleDelSameRef: 37
Lifted OkRefAltSwitchIndelFlank: 389
Lifted RefNewAlleleIndelNoSwitch: 380
Lifted RefNewAlleleInsRefChanged: 121
Failed RefNewAlleleInsRefChanged: 192
Lifted OkRefAltSwitchDelToIns: 1
Failed OkRefAltSwitchDelToIns: 227
Failed INFO/AF: 167
```

5) SNPs – GRCh38 to T2T

```
Genozip
Showing counts of o$TATUS (did_i=17). Total items=4082903 Number of
categories=7
OkRefSameSNP        2288044     56.04%
OkRefAltSwitchSNP       1705586     41.77%
RefNotMappedInChain     61771 1.51%
OkRefSameSNPRev  22266 0.55%
RefNewAlleleSNP  1934  0.05%
OkNewRefSNP      1869  0.05%
RefMultiAltSwitchSNP  1433  0.04%

CrossMap
data=snp primary=38 luft=t2t tool=CrossMap
Lifted OkRefSameSNP: 2288044
Failed OkRefAltSwitchSNP: 1705586
Failed RefNotMappedInChain: 61771
      Lifted OkRefSameSNPRev: 22266
Lifted RefNewAlleleSNP: 1934
Lifted OkNewRefSNP: 1869
Lifted RefMultiAltSwitchSNP: 1433

LiftoverVcf
data=snp primary=38 luft=t2t tool=gatk
Lifted OkRefSameSNP: 2288044
Lifted OkRefAltSwitchSNP: 1705586
Failed RefNotMappedInChain: 61771
Lifted OkRefSameSNPRev: 22266
Failed RefNewAlleleSNP: 1934
Failed OkNewRefSNP: 1869
Failed RefMultiAltSwitchSNP: 1433
```

6) ClinVar – GRCh38 to T2T

```
Genozip
Showing counts of o$TATUS (did_i=17). Total items=969486 Number of
categories=22
OkRefSameSNP	857612	88.46%
OkRefSameIndel	86095	8.88%
OkRefAltSwitchSNP	11055	1.14%
OkRefSameNotLeftAnc	5972	0.62%
OkRefSameSNPRev	2058	0.21%
RefNewAlleleInsSameRef	1071	0.11%
RefSplitInChain	1003	0.10%
RefNotMappedInChain	859	0.09%
OkRefAltSwitchIndelRpts	734	0.08%
RefNewAlleleSNP	732	0.08%
RefNewAlleleDelRefChanged	520	0.05%
RefNewAlleleNotLeftAnc	457	0.05%
OkRefAltSwitchWithGap	432	0.04%
RefNewAlleleDelSameRef	340	0.04%
OkRefAltSwitchIndelFlank	152	0.02%
RefNewAlleleInsRefChanged	111	0.01%
OkRefSameDelRev	89	0.01%
OkRefAltSwitchDelToIns	53	0.01%
OkRefAltSwitchNotLeftAnc	50	0.01%
OkRefSameInsRev	48	0.00%
RefNewAlleleIndelNoSwitch	42	0.00%
ChromNotInPrimReference	1	0.00%

CrossMap
data=clinvar primary=38 luft=t2t tool=CrossMap
Lifted OkRefSameSNP: 857612
Lifted OkRefSameIndel: 86095
Failed OkRefAltSwitchSNP: 11055
Lifted OkRefSameNotLeftAnc: 5972
Lifted OkRefSameSNPRev: 2058
Lifted RefNewAlleleInsSameRef: 1071
Lifted RefSplitInChain: 106
Failed RefSplitInChain: 897
Lifted RefNotMappedInChain: 68
Failed RefNotMappedInChain: 791
Lifted OkRefAltSwitchIndelRpts: 734
Lifted RefNewAlleleSNP: 732
Lifted RefNewAlleleDelRefChanged: 520
Lifted RefNewAlleleNotLeftAnc: 456
Failed RefNewAlleleNotLeftAnc: 1
Lifted OkRefAltSwitchWithGap: 1
Failed OkRefAltSwitchWithGap: 431
Lifted RefNewAlleleDelSameRef: 340
Lifted OkRefAltSwitchIndelFlank: 152
Lifted RefNewAlleleInsRefChanged: 111
Lifted OkRefSameDelRev: 89
Lifted OkRefAltSwitchDelToIns: 53
Lifted OkRefAltSwitchNotLeftAnc: 16
Failed OkRefAltSwitchNotLeftAnc: 34
```

```
Lifted OkRefSameInsRev: 48
Lifted RefNewAlleleIndelNoSwitch: 42
Failed ChromNotInPrimReference: 1

LiftoverVcf
data=clinvar primary=38 luft=t2t tool=gatk
Lifted OkRefSameSNP: 857612
Lifted OkRefSameIndel: 86095
Lifted OkRefAltSwitchSNP: 11055
Lifted OkRefSameNotLeftAnc: 5972
Lifted OkRefSameSNPRev: 2058
Lifted RefNewAlleleInsSameRef: 1071
Failed RefSplitInChain: 1003
Failed RefNotMappedInChain: 859
Lifted OkRefAltSwitchIndelRpts: 734
Failed RefNewAlleleSNP: 732
Failed RefNewAlleleDelRefChanged: 520
Failed RefNewAlleleNotLeftAnc: 457
Failed OkRefAltSwitchWithGap: 432
Lifted RefNewAlleleDelSameRef: 340
Lifted OkRefAltSwitchIndelFlank: 152
Lifted RefNewAlleleInsRefChanged: 2
Failed RefNewAlleleInsRefChanged: 109
Lifted OkRefSameDelRev: 87
Failed OkRefSameDelRev: 2
Failed OkRefAltSwitchDelToIns: 53
Failed OkRefAltSwitchNotLeftAnc: 50
Lifted OkRefSameInsRev: 48
Lifted RefNewAlleleIndelNoSwitch: 42
Failed ChromNotInPrimReference: 1
```
